# Long-term evolution of proliferating cells using the eVOLVER platform

**DOI:** 10.1101/2023.03.28.534552

**Authors:** Daniel García-Ruano, Akanksha Jain, Zachary J Heins, Brandon G Wong, Ezira Yimer Wolle, Ahmad S Khalil, Damien Coudreuse

## Abstract

Experimental evolution using fast-growing unicellular organisms is a unique strategy for deciphering the principles and mechanisms underlying evolutionary processes as well as the architecture and wiring of basic biological functions. Over the past decade, this approach has benefited from the development of powerful systems for the continuous control of the growth of independently evolving cultures. While the first devices compatible with multiplexed experimental evolution remained challenging to implement and required constant user intervention, the recently-developed eVOLVER framework represents a fully automated closed-loop system for laboratory evolution assays. However, it remained difficult to maintain and compare parallel evolving cultures in tightly controlled environments over long periods of time using eVOLVER. Furthermore, a number of tools were lacking to cope with the various issues that inevitably occur when conducting such long-term assays. Here we present a significant upgrade of the eVOLVER framework, providing major modifications of the experimental methodology, hardware and software as well as a new standalone protocol. Altogether, these adaptations and improvements make the eVOLVER a versatile and unparalleled setup for long-term experimental evolution.

## INTRODUCTION

Over the last decades, experimental evolution approaches have gained great interest not only for the field of evolutionary biology, but also for investigating fundamental questions in cell biology. Evolving cells in controlled conditions provides an unprecedented entry point for unraveling novel biological mechanisms and exploring unanticipated areas of cellular function (Anderson et al. 2004; Blount et al. 2008; Bull 2010; Torres et al. 2010; Gresham and Dunham 2014; Kryazhimskiy et al. 2014; Hope et al. 2017; Fumasoni and Murray 2021; Pavani et al. 2021). It is also a unique method for studying long-term genome dynamics and reorganization (Yona et al. 2012; Lang et al. 2013; Maddamsetti et al. 2015; Vázquez-García et al. 2017). Moreover, laboratory evolution is a key component of biological engineering projects and applied research, for instance for improving the yield of *in vivo* cellular factories (Dragosits and Mattanovich 2013; Mundhada et al. 2017; Wang et al. 2023).

While the sequential dilution of batch cultures paved the way for deciphering the evolutionary biology of unicellular organisms from an experimental perspective (Lenski and Travisano 1994; Elena and Lenski 2003), the recent emergence of new methods and devices allows for performing such assays in a more controlled manner (Klein et al. 2013; Miller et al. 2013; Toprak et al. 2013; Takahashi et al. 2015; Callens et al. 2017; Wong et al. 2018). Such systems were developed to simultaneously evolve several independent populations in small culture volumes while remaining cost-effective compared to standard larger chemostats. One such device, the “ministat” array, represented a major breakthrough for conducting experimental evolution and was successfully implemented for both budding and fission yeast (Miller et al. 2013; Callens et al. 2017). However, this approach remains difficult to sustain in turbidostat mode and is very sensitive to small changes in the setup (*e.g.* needle height, pump tubing), thus requiring constant monitoring by the user. Furthermore, while the simplicity of the ministats is a major advantage of the system, the absence of automatization, real-time recording of culture optical densities (OD) and feedback on the dilution rate makes it challenging to maintain stable cultures.

More recently, the eVOLVER framework was designed as the first open-source and automated system for experimental evolution (Wong et al. 2018; Heins et al. 2019). This device measures the OD of each evolving population, triggering controlled dilutions in user-defined density ranges (Fig. 1A). Furthermore, each vial can be run with a different medium, temperature, and culture agitation rate, providing an unprecedented level of versatility. However, when performing long-term assays over several months, we encountered a number of issues that made it difficult to maintain consistent culture conditions and obtain robust datasets. In particular, we observed differences in the accuracy of the OD measurements between positions as well as alterations in OD readings upon changing a culture vial in a given sleeve, a situation that occurs in the case of a dilution problem or vial contamination. This represented a major drawback for comparing the growth rates and evolution dynamics between experimental replicates or between populations of genetically distinct strains. We therefore set out to improve the eVOLVER platform for the laboratory evolution of exponentially growing cell cultures. Here we describe a number of critical modifications of the eVOLVER system, from its calibration procedure to its hardware, and provide a set of newly-developed methods that are essential for performing long-term evolution assays. We discuss the rationale and principles of the major improvements that we have implemented and demonstrate their advantages using yeast cells as a proof-of-concept. We also include a comprehensive and standalone Materials and Methods section in the Supplementary Information, together with all the scripts necessary to conduct evolution experiments using unicellular micro-organisms. These changes to the original system can be applied to both commercial setups and home-made eVOLVERs. Altogether, our modified system and protocols can be used for long-term assays by any research group without prior experience or know-how in experimental evolution.

**Figure 1.**
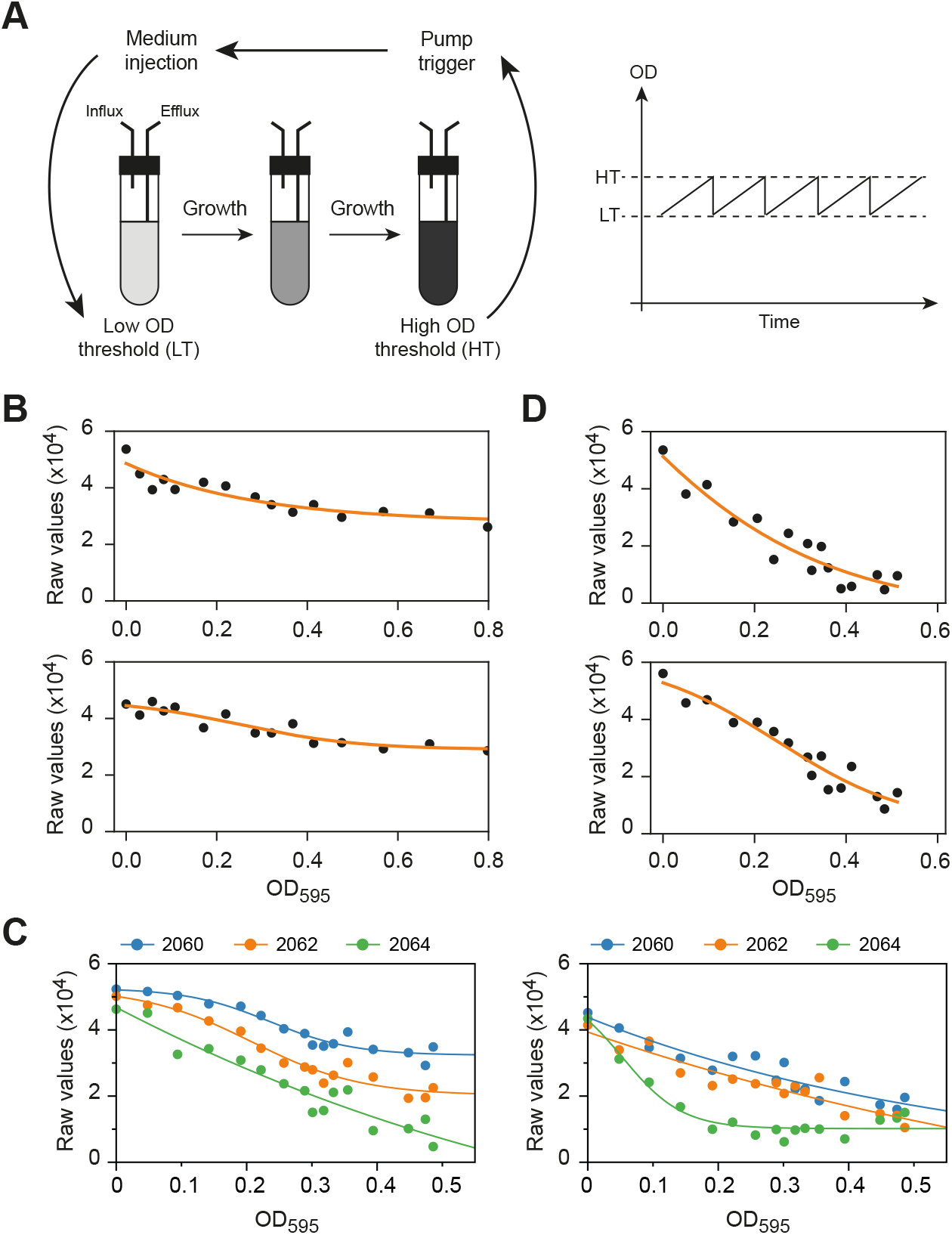
**A.** Principle of the eVOLVER system. Left panel: cells are grown in dedicated culture vials, whose optical densities are constantly monitored by eVOLVER. When a culture reaches a user-defined high OD threshold (HT), the influx pump that is connected to the vial is automatically triggered, allowing for the injection of medium until the culture OD reaches a user-defined low OD threshold (LT). A constant culture volume is maintained due to the concomitant operation of the efflux pump during the dilution process. Right panel: the eVOLVER system allows for maintaining continuous cultures that oscillate between two OD thresholds. Using a narrow range of ODs allows a turbidostat-like operation of the system. **B.** OD calibration curves of two representative eVOLVER sleeves generated using the original calibration protocol (Wong et al. 2018; Heins et al. 2019). **C.** Using the updated eVOLVER hardware, the LED power can be adjusted for each sleeve to improve the dynamic range of the OD calibration curves. The curves of two representative sleeves with three different LED powers are shown. These curves were obtained using the original calibration method. **D.** OD calibration curves of two representative eVOLVER sleeves after LED optimization, using the original calibration method. All experiments were performed using wild-type fission yeast cells grown in minimal medium (EMM) and subsequently diluted in PBS to prevent growth during the calibration procedure.

## RESULTS AND DISCUSSION

### Improving the quality and reproducibility of optical density measurements

#### Optimizing LED power

Multiplexed experimental evolution assays using the eVOLVER platform require accurate measurement of population optical densities (OD) in each of the individual vials. However, the system does not directly record OD but rather converts light scattering readings into OD using a user-established calibration curve (Wong et al. 2018; Heins et al. 2019). The method that is currently integrated in the eVOLVER framework for calibrating OD measurements relies on the permutation of 16 different non-growing cultures of pre-determined OD in each of the 16 sleeves of eVOLVER and records the corresponding light scattering values. The calibration curve is then stored internally in the system and used to determine the OD of evolving cultures during and between experiments. While relatively simple, this method produces calibration curves that are established as a fit of heterogenous data points, and there are significant differences in the quality and dynamic range of the calibration curves between sleeves (Fig. 1B). In particular, relatively flat curves lead to highly imprecise and noise-sensitive measurements. To improve the dynamic range of the curves, a hardware upgrade was recently introduced to optimize the power of the emitting LED of each sleeve. To take advantage of this, a set of successive standard calibrations of the OD readings must be performed using different LED outputs, and the most appropriate values are then determined for each vial based on the obtained set of curves (Fig. 1C). To facilitate this procedure, we are providing the script *Cal_LEDpower* that not only allows for setting the LED power during this initial step, but also for reconfiguring all sleeves with the final set of selected values to be used for subsequent experiments (Supplementary Information). Note that for exponentially growing yeast cells, the target OD range (∼0.2 to 0.5) only requires the use of the 135 degrees photodiode of the machine. Altogether, this initial optimization is essential for the quality of the measurements and needs to be performed only once, although any modification of an eVOLVER sleeve (*e.g*. re-positioning of the LED or photodiode) requires a new LED tuning.

#### A new OD calibration method for improved accuracy

Despite the improved dynamic range following the LED power optimization, there is still significant noise in the calibration measurements. The scattering *vs*. OD curves thus remain imprecise fits (Fig. 1D). This has implications for the quality of the OD determination, resulting in major differences in OD values when comparing the readings from eVOLVER with those from a standard and accurate spectrophotometer (Fig. 2A). This is a critical issue when performing experimental evolution assays: such differences make it difficult to reliably determine the population doubling time, influencing the frequency of culture dilution, which is set by OD thresholds, and preventing the robust comparison of population behaviors between vials.

**Figure 2.**
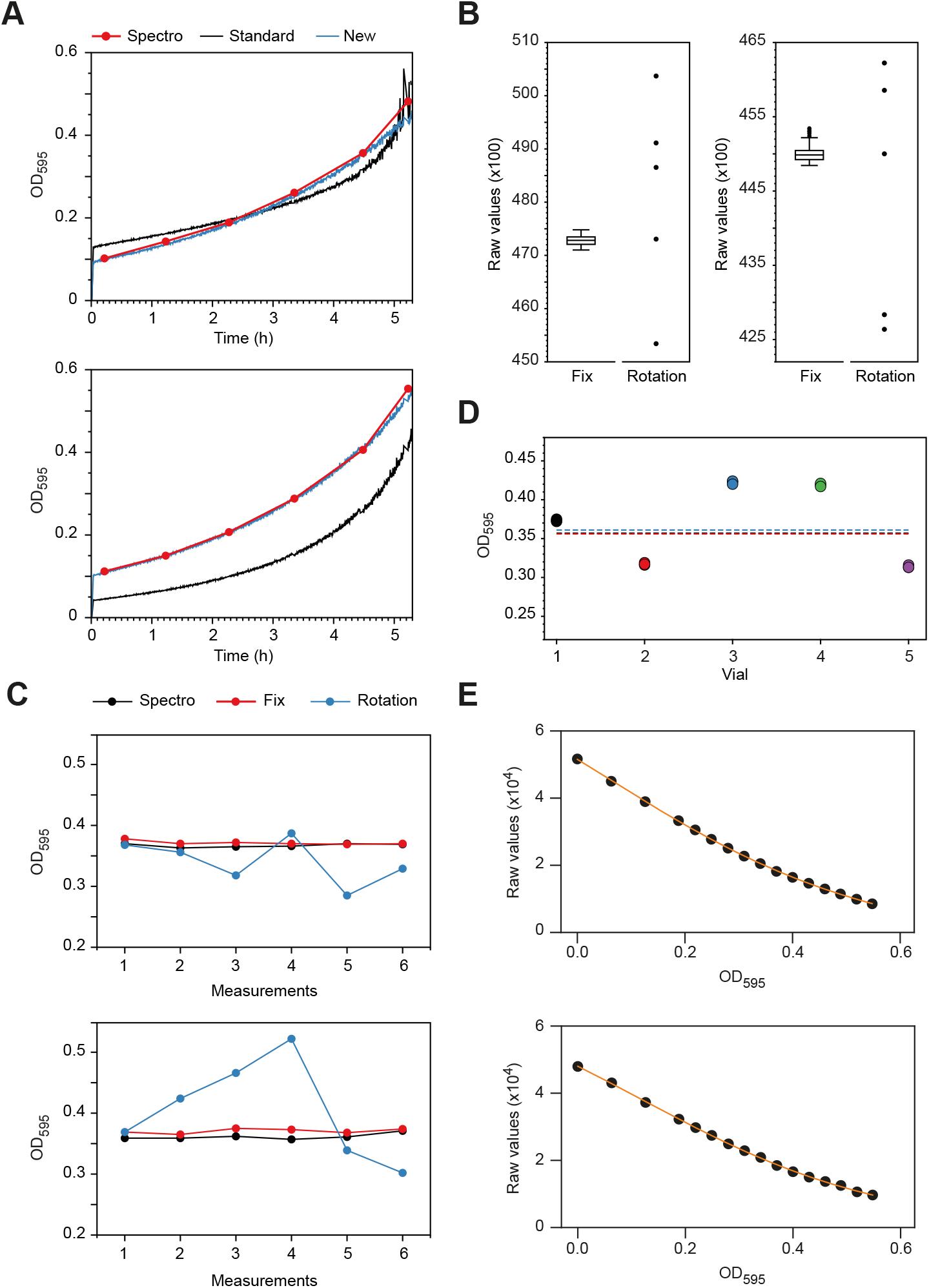
**A.** OD measurements of growing cultures in eVOLVER. Two representative sleeves are shown. At T=0, wild-type fission yeast cells were inoculated in minimal medium (EMM), and the OD of the populations were monitored using 1) culture sampling and OD determination in a spectrophotometer (Spectro), 2) the original calibration method after LED optimization (Standard) and 3) the new calibration method that we developed, after LED optimization (New). For this experiment, no automated dilution was applied. **B.** Changes in the position of the glass vial within a smart sleeve strongly impact the accuracy of the raw measurements. The data for two representative sleeves are shown. First, the raw values of the light scattering using glass vials filled with PBS were measured continuously for 2h (447 datapoints – Fix; box and whiskers plot indicating the minimum, Q1, median, Q3 and maximum, with outliers determined by 1.5 IQR). Subsequently, 5 successive measurements were made, rotating the vials a few degrees between each acquisition (Rotation). The variability that results from vial rotation is significantly higher than the noise of the measurements observed with a fixed vial. **C.** Changes in the position of the glass vial within a smart sleeve strongly impact the quality of the OD determination. Vials with cells at an OD of ∼0.35 were inserted into the sleeves. The OD of the cultures as determined by eVOLVER were recorded for 5 consecutive time points (1 min between points) first without moving the vials and then with a rotation of a few degrees of the vials between time points. As a control, the ODs of the populations were measured 5 consecutive times using a standard spectrophotometer. The data for two representative sleeves are shown. **D.** Impact of the glass vial on the reproducibility of the OD measurements. Cells were grown to an OD of ∼0.35 as measured three times using a standard spectrophotometer at the beginning of the experiment (dashed lines). The culture was then split in 5 eVOLVER glass vials and the OD of each vial was determined within the same sleeve (3 consecutive points at 1 min intervals for each vial). **E.** OD calibration curves of two representative eVOLVER sleeves using optimized LED power and the new calibration method. *B-E:* All experiments were performed using wild-type fission yeast cells grown in minimal medium (EMM) and subsequently diluted in PBS to prevent growth during the calibration procedure.

We hypothesized that this noise in the calibration measurements and the subsequent inaccuracy in OD determination during an evolution experiment may result from the impact of the quality of the glass vials on the recording of the light scattering. Indeed, due to the permutation protocol mentioned above, each data point of the calibration curve is acquired using a different vial, and evolution assays are then performed using yet a different set of vials. This parameter is neglected in the currently available protocol for setting up an eVOLVER assay. To test this possibility, we first assessed whether simply rotating a vial inside a sleeve by a few degrees changes the scattering measurements (Fig. 2B). Strikingly, we observe variations in the data that are significantly higher than the basal noise of the measurements. Importantly, when using a yeast cell culture of known OD, we found that rotation of the vial inside the sleeve leads to significant inaccuracies in the OD value calculated by eVOLVER (Fig. 2C). Furthermore, comparison of the ODs of the same yeast culture in different glass vials measured in a single eVOLVER sleeve revealed major variability in the values (Fig. 2D).

In order to improve the quality of both the calibration curves and the OD determination during an eVOLVER experiment, we established a new calibration protocol that eliminates the impact of the glass vials on the recording of the light scattering (Supplementary Information). This method is based on sets of calibration measurements that are made using a single, fixed vial in each eVOLVER sleeve. Thus, while the integrated software for calibrating the machine is still used, the vial permutation between sleeves is replaced by an initial measurement using PBS (phosphate buffered saline, which prevents cells from growing during the calibration) followed by the inoculation of increasing numbers of cells from a stock culture in each vial for each datapoint. Once the last calibration measurement is performed, the ODs of each culture are measured and averaged using a standard spectrophotometer. This allows for the initial OD of the stock culture to be back-calculated based on the corresponding dilution factor (note that when conducting a more complex experiment using different stock cultures for specific subsets of sleeves, distinct averages must be made to back-calculate the ODs of each individual stock culture). This procedure compensates for potential inaccuracies in the initial determination of the stock culture OD, which is particularly prone to errors. Indeed, this culture is highly concentrated prior to starting the calibration in order to limit the impact of the inoculation steps on the total volume of the cultures in the eVOLVER vials. Using this new value for the stock population, the actual ODs of the cultures at each calibration point are re-calculated and this dataset is then used to correct the calibration data (see below). We are providing the calculation form *Cal_Inoculation* that helps with determining the successive inoculation steps and resulting ODs taking into account the OD of the stock culture (Supplementary Information).

However, when applying our calibration method, OD calibration curves cannot be generated using the standard eVOLVER software. Indeed, the original calibration method relies on 1) the rotational permutation of the vials between sleeves, which implies that the order of the measurements is different between sleeves, and 2) the non-corrected OD values at each point determined prior to back-calculation of the stock culture OD. Thus, to obtain the appropriate calibration curves, the order of the measurements and calibration ODs must be corrected in the file generated by the eVOLVER software. This is achieved using the newly-provided script *Cal_correction* (Supplementary Information). Importantly, our method results in significant improvement of the calibration curves (Fig. 2E), leading to highly accurate determination of the culture densities in eVOLVER vials (Fig. 2A).

To perform robust laboratory evolution assays over long periods of time, a calibration using this new method should be performed prior to each experiment. This contrasts with the standard protocol, in which a single round of calibration of the sleeves is used for multiple independent assays. In addition, the same set of vials, fixed in a given position in each sleeve, should be used for both the calibration step and the ensuing evolution assay. This further improves the quality of the experimental measurements. To this end, vials are flushed after the calibration using fresh medium and the eVOLVER pump system, and cells are then newly inoculated according to the design of the experiment. It is critical that the same strain is used for both the calibration and the experiment so that the limited number of cells remaining from the calibration procedure do not alter the evolution assay. Thus, when different strains are evolved in parallel, each vial must be calibrated using the strain it will subsequently host. This also contributes to the quality of the measurements, as the light scattering of a cell culture is affected by various factors, including cell shape and size. Finally, in contrast to the currently published approach, a given calibration is only used for the accompanying experiment. We are therefore providing the script *Cal_delete* for removing obsolete calibrations (Supplementary Information).

Altogether, while this calibration strategy takes more time to carry out, it substantially improves the quality of the OD measurements, which is critical for maintaining the quality of experiments that last for several months. Our method therefore represents a clear advance for long-term assays, enhancing the control of population densities and allowing for accurate estimation of their growth rates. This makes it possible to take full advantage of the possibility of evolving multiple replicates or comparing strains and conditions within a single run.

### Alteration of the temperature calibration range

Calibration of the temperature for each eVOLVER sleeve is a key step when setting up an eVOLVER machine. Indeed, the temperature of an eVOLVER culture is determined based on the temperature measured at the surface of the aluminum cylinder that makes up the sleeve. The relationship between these two temperatures must therefore be established for each sleeve. The eVOLVER software integrates a temperature calibration procedure based on three measurements: room temperature (RT), RT + ∼10 °C and RT + ∼20 °C. However, when this protocol is performed at a RT greater than 20 °C, the system often fails to reach the RT + ∼20 °C temperature. We noticed that the lack of stability of the 3^rd^ temperature point can lead to inaccuracy in the working temperatures that are then set for the evolution assay. Furthermore, depending on the RT, the temperature of the third point is generally outside of the physiological range. To eliminate this source of variability, we have modified the eVOLVER software source code and compiled a new application *eVOLVER_V2*, which uses RT, RT + ∼6 °C and RT + ∼12 °C (Supplementary Information). This modification generates accurate temperature calibration curves and improves the reliability of the culture temperature during an evolution assay. While the calibration target values were established based on standard growth temperatures of fission and budding yeast cells (25 to 32 °C), this range can be modified by the user to generate a new compilation of the eVOLVER application (Supplementary Information). We noted that despite carefully performing this calibration, a limited temperature offset may persist between the temperature determined by eVOLVER and the temperature measured using a highly accurate sensor directly immersed in the culture (0.15 °C lower, on average, than the target temperature in our experiments; data not shown). While this offset is largely negligible, it can be determined for each vial and used to correct the target temperature when an experiment requires an extremely accurate control of the growth temperature.

### Long-term experimental evolution assay

#### OD calculation during an evolution experiment

An eVOLVER assay begins with the blank measurements of vials that contain only growth medium. The raw_blank_ value (light scattering value) and the calibration curve (OD=f(raw)) of the sleeve then allow for determining an intermediate OD value that we refer to as OD_blank_. Cells are then inoculated and at a time T1, the OD of the culture is determined as follows. A raw_T1_ value is first measured, and an intermediate OD_interT1_ is established using the calibration curve. The final absolute OD is calculated as OD_interT1_ - OD_blank_. However, using the original eVOLVER code and calibration procedure, this approach can introduce significant errors in OD determination, as both OD_blank_ and OD_interT1_ are calculated from the raw measurements and the inaccurate, noisy calibration curves (see above).

While the new calibration and experimental methods using fixed glass vials generate high-quality calibration curves, we reasoned that the use of the raw data for determining the difference in measurements between the culture and the blank is a more reliable strategy to calculate the OD of a given population. First, we establish a difference in raw values Δraw_T1_ = raw_T1_-raw_blank_. As these two raw values are measured from the same fixed glass vial, Δraw_T1_ results solely from changes in cell culture density. Nevertheless, by transposing this difference to the calibration curve, we compensate for any small variations that may result from limited movements of the glass vial. Thus, the OD of the culture is calculated using the calibration curve and a light scattering value raw_final_ = raw_0_cal_ + Δraw_T1_, with raw_0_cal_ being the raw value of the first measurement obtained during the calibration, using PBS.

It is important to note that these methods can only be applied when there is no major difference between the light scattering of the medium and that of PBS. We have validated this through comparing the raw light scattering data from eVOLVER as well as the OD determined by a spectrophotometer for different media that are commonly used to grow fission yeast cells (Table 1). We have now implemented the OD calculation based on Δraw_T1_ in the eVOLVER code that we provide (Supplementary Information).

**Table 1.** Light scattering measurements of different growth media. Raw values of light scattering were determined for different solutions and fission yeast media using a single fixed glass vial in a single eVOLVER sleeve. For each condition, 5 consecutive raw measurements were recorded (averages and standard deviations are shown) and the differences in average raw values with that of water were calculated. OD_595_ were determined using a standard spectrophotometer, with water used as a blank. The differences in light scattering between the solutions are in the range of the measurement noise (Fig. 2B). PBS is the standard solution for the calibration procedure. EMM: minimal medium. EMM6S: supplemented minimal medium. YE4S: supplemented yeast extract-based rich medium.

|  | **Water** | **PBS** | **EMM** | **EMM6S** | **YE4S** |
| --- | --- | --- | --- | --- | --- |
| **Raw value** | 41295 ± 33 | 41115 ± 30 | 41652 ± 54 | 41362 ± 79 | 41457 ± 66 |
| **Difference Raw** | 0 | -180 | 358 | 68 | 162 |
| **OD<sub>595</sub>** | Blank | 0.00 | 0.00 | 0.00 | 0.03 |

#### Hardware improvements for long-term evolution assays

As discussed above, changes in the exact position of the glass vials in the smart sleeves are problematic for both the calibration and the evolution assay, as they have a strong impact on the OD determination. We have therefore designed new 2-part caps that prevent vial rotation while providing the user with all the necessary ports for connecting the vials to the pump system and for sampling the cultures (Fig. 3A). These caps consist of a base ring that is attached to the sleeve, combined with a top complementary part that is fixed to the glass vial. CAD files are provided for 3D-printing of these caps, which can be produced from autoclavable materials (Supplementary Information).

**Figure 3.**
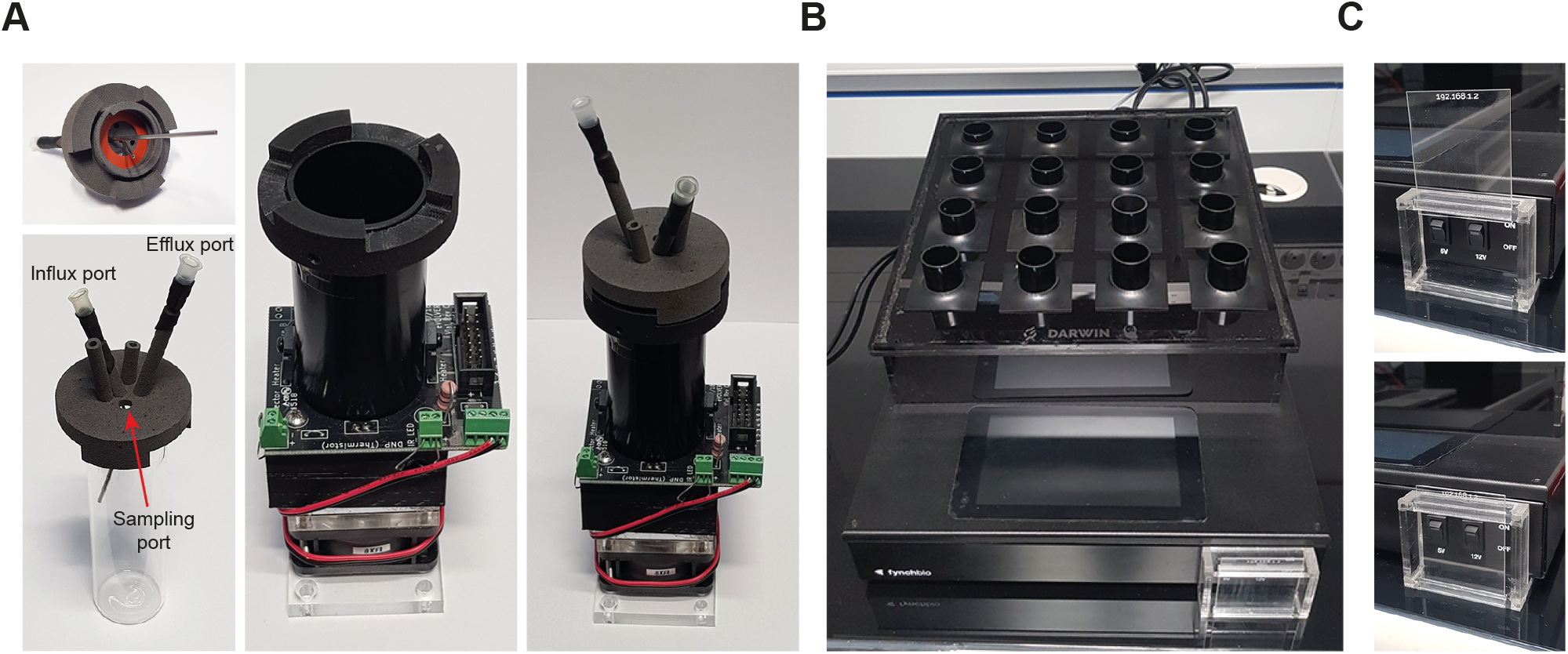
**A.** Newly designed vial caps that allow for fixing the position of the vials in the sleeves while preventing leaked medium from flooding the sleeves. The needles are maintained in position using standard heat-shrink tubing. The cap integrates two additional ports that can be used for more complex assays. **B.** eVOLVER enclosure designed to protect the main body of the eVOLVER machine from leakage of liquid medium. The casing is tilted to facilitate the efficient evacuation of liquid medium. **C.** Casing designed to prevent accidental switching off of the eVOLVER system during an experiment.

One of the major problems encountered when performing long-term experimental evolution assays using eVOLVER is the possibility of medium leaks from the vials. Most of the time, these result from alteration of the effluent pump operation or clogging of the effluent line. As it is difficult to fully avoid such leaks, we have designed additional hardware that protects both the sleeves and main body of the eVOLVER from being exposed to liquid medium. First, the new caps described above have a diameter that is larger than that of the sleeves themselves (Fig. 3A). This prevents medium that may leak from the connectors or sampling port from flooding the sleeve and damaging its electronic board. Silicone rings are used to seal the cap to the vials. In addition, we have fabricated a protective casing for the eVOLVER: combined with silicon pads, it protects the main eVOLVER hardware from being damaged (Fig. 3B). Altogether, these improvements limit the risk of aborting an ongoing long-term evolution assay.

Finally, over the course of an evolution experiment, the user regularly manipulates the eVOLVER system and associated tubing. We noticed that the positioning of the eVOLVER main switch makes it possible to accidentally turn the machine off, entirely stopping the ongoing experiment. We are therefore using a casing system that protects the power switch (Fig. 3C). All of these parts can be fabricated using a standard CO_2_ laser cutter, using the accompanying vector files (Supplementary Information).

#### Restarting an evolution assay from a problematic vial

One of the most critical aspects of running a long-term experimental evolution assay using eVOLVER is to have the possibility to restart a culture when a specific vial fails, generally due to contamination or medium leakage. Importantly, this must be achieved without interfering with the other vials. First, this requires the sampling and freezing of all ongoing cultures at regular and frequent intervals, making it possible to resume an experiment from a well-defined time point. Second, the overall experiment must be paused, a new vial inserted in the specific failed position and inoculated with cells, followed by resumption of the experiment. However, this procedure is not straightforward: indeed, switching a failed vial for a new one with a freshly thawed culture has an impact on the OD determination due to the change of glass vial, as discussed above.

We have therefore established a robust method for dealing with this frequently occurring and crucial issue. First, a new vial containing only growth medium is inserted. It is then manually rotated to reach an OD close to zero, as determined by eVOLVER. To facilitate this, we use the *VialReplace_Equilibrate* script, which displays the OD determined by eVOLVER in real-time. Note that the OD indicated in the eVOLVER application or on the touch screen of the machine cannot be used as it takes into account the calibration blank rather than that of the growth medium. If a value of zero is reached or when the difference with the blank that was determined at the start of the assay is considered by the user as non-significant for the assay, the cells can be inoculated and the experiment resumed.

However, it is often difficult to reach an OD that is sufficiently close to zero, due to the high variability between glass vials. In this case, the calibration curve of only the corresponding position must be corrected to integrate the vial change. This is achieved through resetting the calibration blank (Fig. 4), using the *VialReplace_Blank* script. A complete description of this critical protocol is provided in the Supplementary Information. This method allows the user to maintain a high degree of accuracy in OD measurements and doubling time determinations despite the change in glass vial. Notably, this replacement procedure can also be used if an experiment is designed to integrate additional cultures during an ongoing assay. To this end, all vials should be calibrated prior to starting the experiment, and the addition of new vials should simply be treated as the replacement of a failed vial.

**Figure 4.**
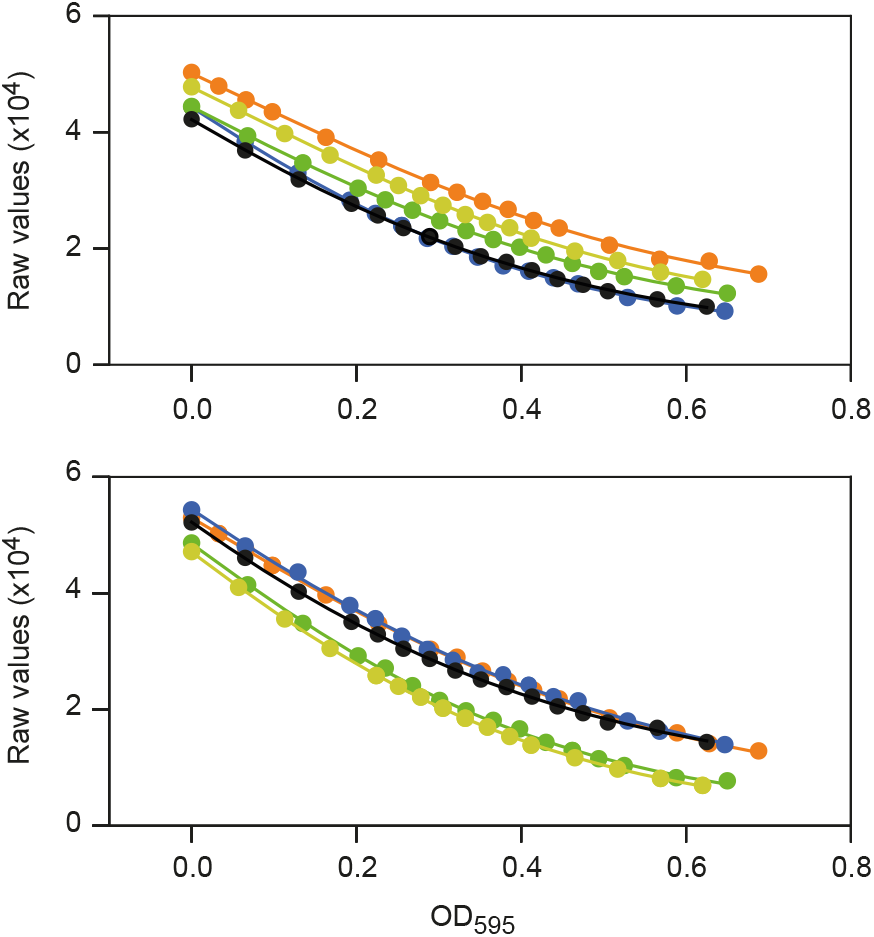
Overlay of distinct calibration curves obtained by performing five consecutive calibrations of a single sleeve using five distinct vials. Data for two representative sleeves are shown. While the overall shape of the curves is mostly determined by the sleeve hardware, the light scattering measurement of the initial calibration sample (PBS) is influenced by the glass vial. Thus, for a given sleeve, the calibration curves that are obtained with different vials are mostly parallel, only differing by a constant offset in the raw scattering data. This makes it possible to use the difference in medium blanks, rather than that in calibration blanks, to correct the calibration curve of a sleeve when the vial needs to be changed during an evolution assay. Experiments were performed using fission yeast cells operating with a minimal cell cycle control network (Coudreuse and Nurse 2010; Banyai et al. 2016) grown in minimal medium (EMM) and subsequently diluted in PBS to prevent growth during the calibration procedure.

Restarting individual cultures from intermediate frozen samples in an ongoing long-term experimental evolution assay is frequently necessary. Here we provide all the tools to deal with this issue while maintaining high quality measurements and data. Together with the other improvements that we have implemented, this enables the user to maintain evolving cultures in reproducible conditions over long periods of time (*e.g*. > 500 generations, Fig. 5)

**Figure 5.**
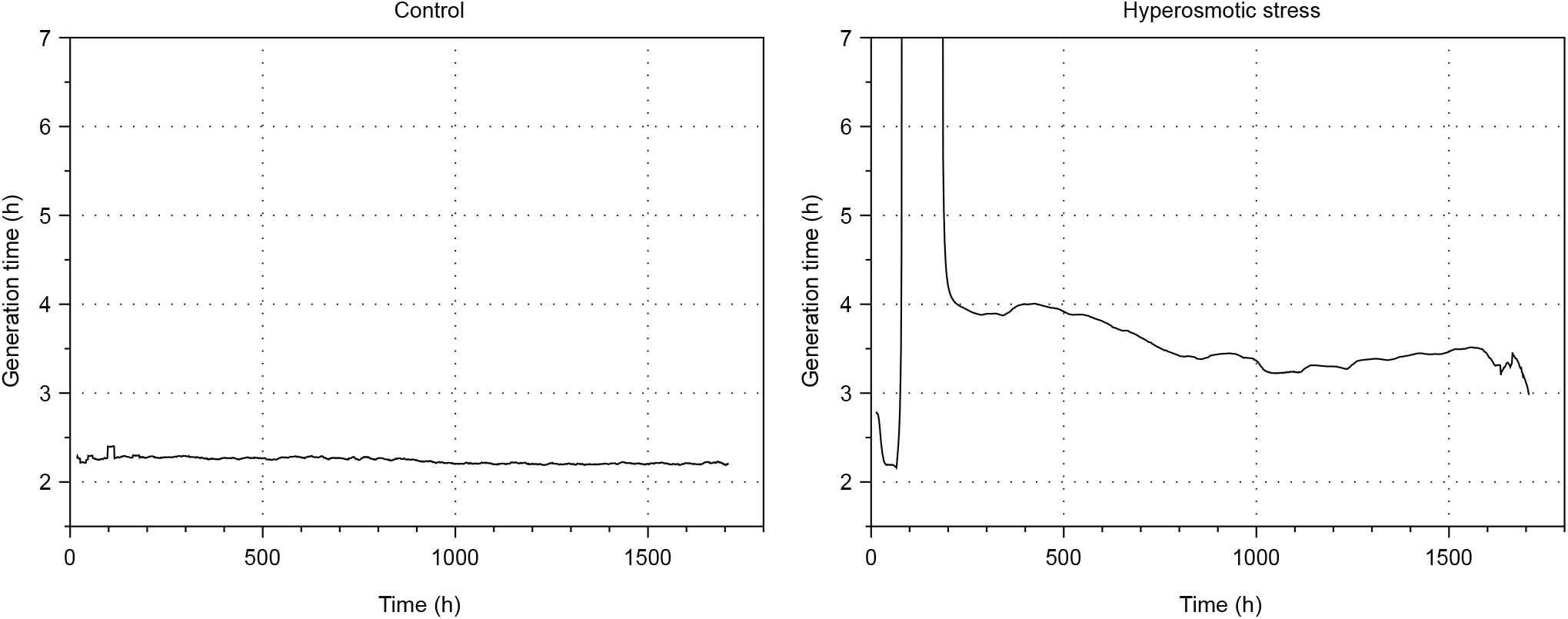
Example of an experimental evolution assay using the improved eVOLVER platform and fission yeast cells growing in the absence of stress (left panel) or exposed to hyperosmotic conditions (right panel). For visualization purposes, the y-axis (generation time) in the stress conditions is limited to 7h, preventing the full display of the generation time curve (maximum generation time: ∼14h). Experiments were performed using fission yeast cells operating with a minimal cell cycle control network (Coudreuse and Nurse 2010) and lacking the flocculin-encoding gene *gsf2*, grown in minimal medium (EMM, left) or in EMM + 1.5 M KCl (right).

### Monitoring a long-term experimental evolution assay

#### An improved interface for monitoring evolving cultures

One of the keys to the success of a long-term experimental evolution assay is the capacity to monitor the critical parameters associated with the experiment. The previous eVOLVER graphic interface allowed the user to separately display the OD measurements of a given culture. We have now made several improvements to this interface (Supplementary Information). First, together with the OD recording, a plot displays the real-time population growth rate, as calculated by the standard eVOLVER code based on the growth curve between two successive dilutions, as well as the average growth rate over time using a sliding window of 10 successive points. Second, in addition to the specific pages dedicated to each culture, the main page now displays the growth rate data of all ongoing cultures, allowing the user to have a rapid overview of the entire assay (Fig. 6A). Third, we have a new *DILUTIONS* page for visualizing the dilution information and media consumption, as well as accessing the tools for the management of the media bottles that are connected to the vials (Fig. 6B). This page includes a setup interface, allowing the user to assign given media bottles to specific sets of vials and subsequently track medium consumption. It also makes it possible to register a change in media bottles, generating a log of all bottle switches over the course of the experiment. Finally, the *DILUTIONS* page displays additional information such as the calculated vial consumptions and the efficiency of the dilution steps (*i.e.* how many dilutions are required to achieve the target; see Supplementary Information).

**Figure 6.**
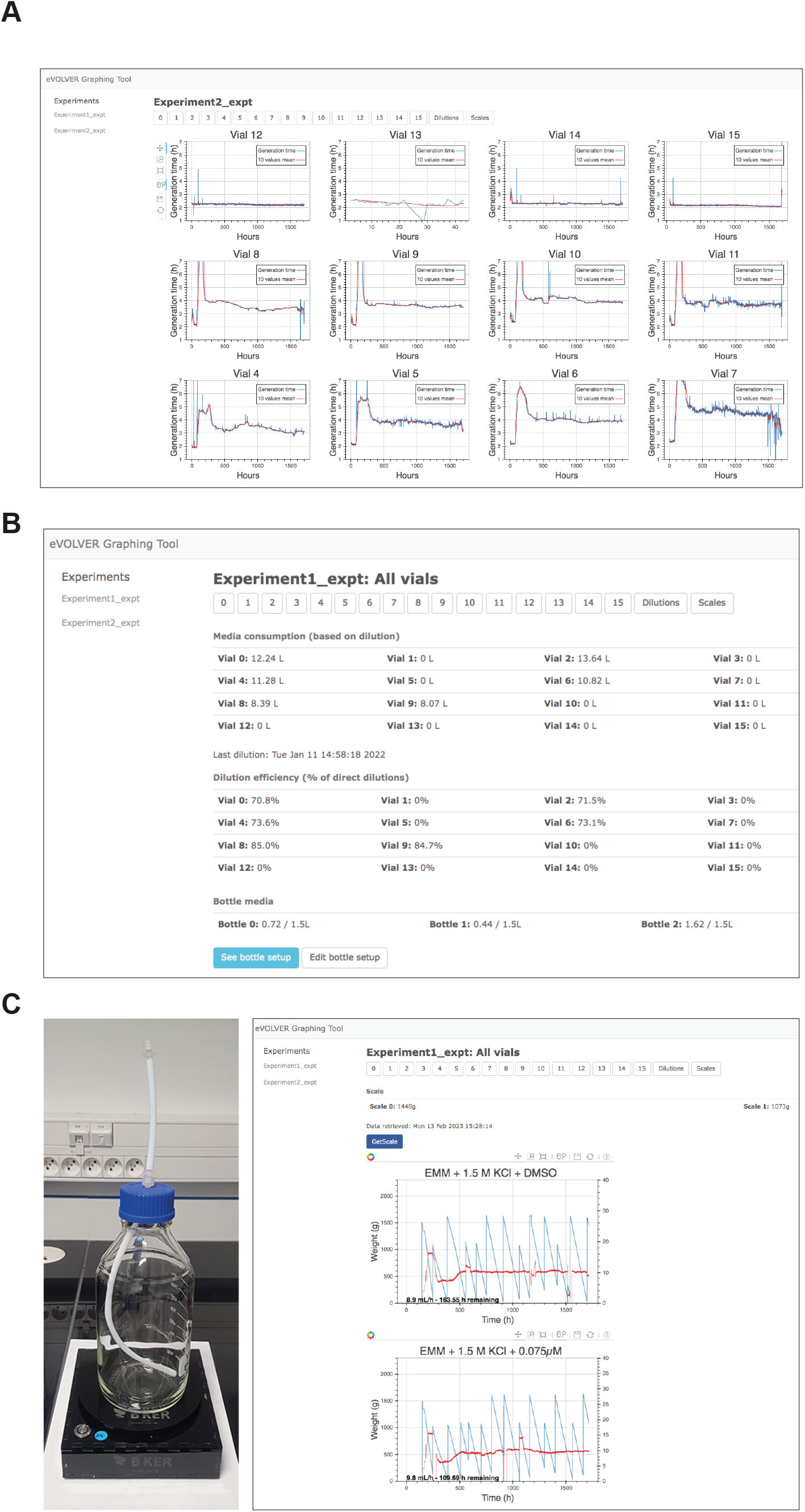
**A.** Screen capture of the main page of the new graphic interface displaying the data for all ongoing cultures (only 12 vials were used in the displayed assay). **B.** Screen capture of the *DILUTIONS* page for dilution monitoring and medium bottle management. Medium consumption is calculated from the start of the experiment. In the displayed assay, not all sleeve positions are used. **C.** Left: Scale dedicated to monitoring the quantity of medium in a feeding bottle. Right: screen capture of the graphic interface dedicated to the scale-based media monitoring system (assay with 2 bottles is displayed as an example).

#### A new set of tools for the surveillance of the eVOLVER system

While performing a long-term evolution assay, it is essential for the user to deal rapidly with potential issues and make sure that all vials remain connected to bottles that contain sufficient amounts of media. To this end, we have integrated a messaging functionality that takes advantage of the open-source Telegram Bots and automatically sends notifications to the user (Supplementary Information). The current version of the messaging functionality that we have implemented alerts the user when an error occurs in the eVOLVER experiment code and blocks the assay (*e.g.* syntax error resulting from modifying the experimental script). It also sends a notification when an experiment is paused for more than 10 min. This is particularly important as an assay is often manually paused when manipulating the machine or sampling the cultures. If the experiment is not resumed, cells grow to saturation, thus impinging on the quality of the results.

In addition, we have developed a tool that monitors medium consumption using individual scales (Fig. 6C) and sends a GSM message when the medium has reached a low limit (Supplementary Information). This utilizes a commercial system (www.bker.io) that we have implemented in the graphic interface. The messaging limits are defined by the user and set on the device’s online portal at the beginning of the experiment, but they can be modified even during an ongoing evolution assay. A *SCALES* page related to this system is available in the improved graphic interface (Fig. 6C). It provides the real-time weight of each bottle as well as graphs of estimated medium consumption. It also includes a calculation of the medium consumption rates and predictions of the time until a bottle change will be required. All the data acquired by the scales are available in the experimental folder. Finally, this system also monitors the power: when connected to the same electrical circuit as eVOLVER, it alerts the user in case of a power cut and complete stoppage of the machine. Altogether, this system is not only useful for alerting when a bottle of media runs empty (note that an unexpectedly high medium consumption can occur in case of a vial failure and thereby affect all vials connected to the same bottle), but it is also a resource that facilitates the planning and anticipation of the different steps that are required to maintain a robust, long-term evolution assay.

## CONCLUSION

The eVOLVER framework is a unique system for automated, multiplexed experimental evolution of unicellular organisms, providing an unprecedented versatility for performing complex experiments. However, a number of improvements were required for conducting reliable long-term assays over hundreds of generations or more and comparing the dynamics of cultures evolved in parallel. These include an increased accuracy and reproducibility in the OD measurements, the establishment of methods for dealing with issues that commonly occur when performing such long experiments, hardware and software upgrades, and access to tools that facilitate the monitoring of ongoing cultures. Here we provide all the methods, scripts and accompanying materials that allow any research group using eVOLVER to exploit these systems to their full potential. This makes the automated laboratory evolution of multiple parallel cultures over long periods of time robust and manageable. Experimental evolution as a research strategy has gained strong interest in a variety of fields, bringing groundbreaking insights into the wiring of cell biological functions. However, its use remains relatively restricted due to the complexities of carrying out such analyses. The methodologies that we have developed will make it possible for a broader community of researchers to take advantage of this this powerful approach, opening the door to new investigations, projects and collaborations.

## MATERIALS AND METHODS

A detailed and comprehensive step-by-step Materials and Methods section is provided in Supplementary Information. It represents a complete standalone methodological guide for performing experimental evolution assays using eVOLVER and the novel methods described above. For fission yeast cell growth and handling, standard methods and media were used (Moreno et al. 1991; Hayles and Nurse 1992). Strains used in this study were PN1 (*972 h-*) and DC684 (*Pcdc13::cdc13Scdc2as::cdc13UTR 𝛥cdc2::KAN 𝛥cig1::HYG 𝛥cig2::KAN 𝛥puc1::HYG 𝛥gsf2::KAN h+*).

## ACKNOWLEDGMENTS

We thank Pei-Yun Jenny Wu and Snezhana Oliferenko for critically reading this manuscript. This work was supported by the Agence Nationale de la Recherche (PRC eVOLve, ANR-18-CD13-0009 to D.C.), the Région Nouvelle Aquitaine (program CHESS, grant agreements 15963520 and 15964420 to D.C.), the National Institutes of Health (NIH) (grants R01AI171100 and R01EB027793 to A.S.K.), and the National Science Foundation (NSF) (grant EF-1921677 to A.S.K.). A.S.K. also acknowledges funding from the Department of Defense (Vannevar Bush Faculty Fellowship N00014-20-1-2825) and the W.M. Keck Foundation. E.Y.W. was supported by an NIH-funded predoctoral training fellowship (T32GM130546). D.G.R. was co-funded by the ANR eVOLve, a grant from the Région Bretagne (ARED) and from the Fondation ARC. A.J. was funded by the ANR eVOLve.

