## Supplementary Information for "Long-term evolution of proliferating cells using the eVOLVER platform"

#### 1. INTRODUCTION

eVOLVER is one of the first open-source, automated and multiplexed systems that was developed for a broad range of applications requiring the continuous culture of micro-organisms, including experimental evolution assays. This device can maintain 16 parallel, independent cultures in a turbidostat mode with minimal user input, making it an ideal and unique tool for performing long-term experiments. The following sections provide a comprehensive and standalone guide for all the steps that are essential to the use of eVOLVER. It combines standard procedures that we previously detailed with a new set of critical improvements that are described in the main manuscript. This will allow researchers without know-how in the use of eVOLVER to rapidly take advantage of this powerful framework for conducting long-term experimental evolution.

### 2. SETTING UP AN EVOLVER

#### 2.1. Hardware

The eVOLVER system is composed of a main unit (*i.e.* motherboard, Arduino microcontrollers, Raspberry Pi, touch screen) that hosts 16 culture sleeves. Each sleeve can be independently controlled for conducting separate assays. Automated dilution of the cultures is ensured by an external rack of peristaltic pumps. This represents the core of the eVOLVER and it should be assembled following manufacturer's instruction.

---

#### 2.2. Software

The eVOLVER hardware elements are coordinated by (1) the integrated Raspberry Pi connected to eVOLVER LCD screen and (2) a computer connected to the eVOLVER by a local network. To setup the network connection between eVOLVER and the computer, refer to this guide.

Next, the complete software and all associated packages needed to control eVOLVER must be installed on the computer. For this, download the following repositories from the SyntheCell GitHub (https://github.com/SyntheCell) and copy them in a folder on the computer (referred to here as the *eVOLVER_computer* folder):

- **DPU**: eVOLVER data processing unit, containing the tools needed to calculate the calibration curves, run an experiment, and monitor eVOLVER data. The DPU requires Python 3 and a set of libraries that are detailed in the *requirements.txt* and *README* files that can be found in the DPU folder.
- **evolver-electron**: source code of the graphical app embedded in the eVOLVER main unit and accessible via the touch screen. The computer version is used to perform the calibration of the system and manually setup a number of parameters (*e.g.* temperature, stirring, pump activation, selection of an existing calibration). If you do not intend to modify the source code, you can directly download the compiled app from the "Releases" section in the repository.
- **evolver-hardware**: repository containing CAD and PDF files of the hardware improvements described in the main manuscript.
- **evolver-utils**: a collection of new python scripts that are notably necessary for optimizing the optical density calibration and replacing vials during an experiment (see main manuscript).

The structure of the *eVOLVER_computer* folder is then as follows:

```
eVOLVER_computer/
├─ dpu/
│  ├─ calibration/
│  ├─ experiments/
│  │  ├─ template/
│  │  ├─ Experiment_1/
│  │  ├─ Experiment_2/
│  ├─ graphing/
│
├─ evolver-electron/
│  ├─ eVOLVER.app
│
├─ evolver-hardware/
│
├─ evolver-utils/
│  ├─ Cal_correction.py
│  ├─ Cal_Inoculation.xlsx
│  ├─ Cal_LEDpower.py
│  ├─ Vial_Blank.py
│  ├─ Vial_Equilibrate.py
```

The template folder inside the DPU contains all scripts needed to run an experiment. Before starting a new experiment, duplicate the template folder and rename it to create a folder that is specific to the new experiment (*e.g.* folders *Experiment_1*, *Experiment_2*). All the data retrieved by eVOLVER during an assay will be saved in the associated folder.

Additionally, you will need to install *ssh* and *scp*, if they are not already available on the eVOLVER computer.

---

##### 2.3. Understanding the experiment folder

The experiment folder, which was duplicated from the template folder (see above), contains all the elements needed to run an eVOLVER experiment. Moreover, after starting an experiment, a new folder recording all the data of the experiment will be created, along with some complementary files. Below is a summary of the files that are within the experiment folder.

| **Tree of the DPU folder** | **Tree of the _*expt* data folder** |
|---|---|
| dpu/ | Experiment_1_expt/ |
| ├─ experiments/ | ├─ |
| │ ├─ template/ | Experiment_1_expt_220317_1745.txt |
| │ ├─ Experiment_1/ | ├─ Experiment_1_expt.pickle |
| │ │ ├─__pycache__/ | ├─ chemo_config/ |
| │ │ ├─scales/ | ├─ evolver.log |
| │ │ ├─Experiment_1_expt/ | ├─ growthrate/ |
| │ │ ├─bottles.txt | ├─ OD/ |
| │ │ ├─**creds.json** | ├─ od_135_raw/ |
| │ │ ├─**custom_script.py** | ├─ ODset/ |
| │ │ ├─**eVOLVER.py** | ├─ pump_log/ |
| │ │ ├─**scales_config.json** | ├─ temp/ |
| │ │ ├─od_cal.json | ├─ temp_config/ |
| │ │ ├─od_raw_zero.json | ├─ temp_raw/ |
| │ │ ├─**pump_cal.txt** | |
| │ │ ├─temp_cal.json | |

The template folder includes the files in **bold**:

- ***creds.json***

  Contains the credentials needed for the Telegram alert bot (see Section 8.4) and the scales system (see Section 6.4).

- ***custom_script.py***

  Contains the main settings and the logic of the turbidostat experiment designed by the user (see Section 6.2).

- ***eVOLVER.py***

  Contains all the logic to start, pause and stop an experiment, as well as to communicate with the eVOLVER machine.

- ***scales_config.json***

  Contains configuration parameters for the scale monitoring tool (see Section 6.4).

- ***pump_cal.txt***

  Contains the pump calibration data for the current experiment (see Section 5.2).

After starting an eVOLVER experiment, several files are created (in green):

- ***od_cal.json***

  Contains the active OD calibration that will be used for the experiment.

- ***od_raw_zero.json***

  Contains the raw light scattering values corresponding to OD=0 of the active calibration. This will be used by eVOLVER to apply the OD blank to the experiment. Additionally, this file can be manually edited to adjust the raw value of each vial at the beginning of an experiment when using an old OD calibration (see Section 6.1).

- ***temp_cal.json***

  Contains the active temperature calibration that will be used for the experiment.

Finally, after configuring the graphing tool, several files are created (in blue):

- ***scales***

  Contains the files *weightXXXX.csv* (where *XXXX* is the scale ID provided by the manufacturer). The scale data are provided in comma-separated value format.

- ***bottles.txt***

  *C*ontains the bottle setup configured in the graphical interface (see Section 6.4) in a tab separated format. The first column corresponds to the bottle ID, the second column to the vials it is connected to, and the other columns detail the dates and times of each bottle changes (odd columns), as well as the volume of each new bottle (even columns). This information will be updated each time a bottle is changed (see Section 7.2).

### 3. ASSEMBLY OF THE COMPONENTS

#### Media bottles

Required parts

- 1L / 2L lab-grade glass bottle
- 2 female luer locks, 4 mm barb
- 1 male luer lock, 4 mm barb
- 1 luer lock cap
- 40 cm of silicone tubing, ID 3.2 mm, OD 6.4 mm

Required equipment

- Drilling machine with 5 mm drilling bit
- Scissors or cutter

*Note:* When using small hydrophobic molecules in the medium, the silicone tubing should be replaced with teflon tubing to limit drug absorption.

1. Drill a hole through the center of the glass bottle cap.
2. Insert the barb of a male luer through the top of the cap, with the luer side facing up.
3. Cut a piece of silicone tubing (see note below) and attach it to the luer lock barb on the inside of the bottle. Note – when using the teflon tubing, attach a 1 cm piece of silicone tubing and a two-way connector between the male luer and the teflon tubing.
4. Cut a 9 cm piece of silicone tubing and insert a female luer lock on each side.
5. Close one end of the part from step 4 using a luer lock cap and attach the assembled part to the female luer lock on the bottle cap (Fig. 1).
6. If needed, use the same tubing with barbed tee splitters to connect the same bottle to several cultures.

*Determining the size of the bottle tubing*

The length of the tubing within the bottle should fit bottle size. The tubing should reach the bottom of the bottle and have a slight bend. Too short tubing does not allow proper medium uptake. Too long tubing makes it difficult to close the bottle cap. We use 24 cm long tubing for 1-liter bottles and 29 cm long tubing for 2-liter bottles.

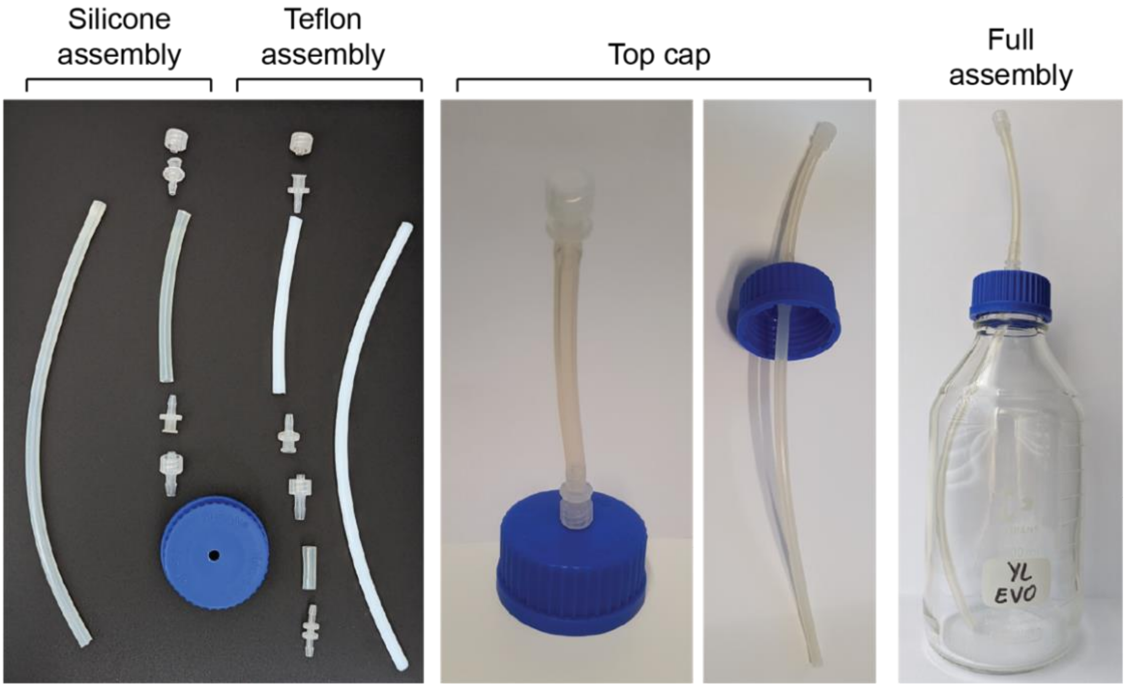

**Figure 1.** Assembly of the media bottles.

---

#### Culture vials

| Required parts | Required equipment |
|---|---|
| • eVOLVER glass vial<br>• Efflux needle (16G, custom length)<br>• Influx needle (18G, 40 mm)<br>• Heat-shrink tubing (6 mm, 2:1)<br>• 3D-printed nylon cap (see Appendix)<br>• Silicon ring (see Appendix)<br>• Magnetic stirring bar | • Heat gun or flame |

1. Determine the length of the efflux needle (see note below).
2. Place a small piece of heat-shrink tubing around the desired port of the top cap.
3. Slide the needle through the port.
4. Hold the heat-shrink tubing and the needle in place so that the heat-shrink tubing overlaps with both the port and the needle. Apply heat to fix the needle.
5. Proceed similarly with the influx needle. The influx needle should be shorter and the cap port should be selected so that it is at a different angle compared to the efflux needle.
6. Place a silicone ring (see Section 8.5) inside of the cap to avoid leaks through the thread. Avoid blocking the sampling port, as this will prevent sampling during the experiment.
7. Put a magnetic stirring bar inside the vial.
8. Screw the assembled cap on a clean and dry glass vial (Fig. 2).

*Determining the length of the efflux needle*

eVOLVER operation is based on a constant culture volume, which is set by placing the efflux needle at a specific height. During the dilution steps, the efflux line will pump out medium until the efflux needle is not in contact with the liquid. This sets the maximum culture volume of the vial. The user must select a culture volume according to the assay and adjust the height of the needle accordingly. This can be achieved by:

- using a needle of specific length
- cutting a standard needle to the appropriate length
- placing the needle in a different cap port
- modifying the height of the needle port before 3D printing the nylon caps

---

**Cap holder**

Required parts

- 3D-printed nylon cap holder (see Appendix)
- M3 x 4 mm headless steel bolts
- EVA foam (2 mm thick)
- Contact glue

Required equipment

- Threader (M3)
- Allen key for the M3 bolts
- Scissors or cutter

1. Thread the three lateral holes in the 3D-printed holder.
2. Cut three 4 mm x 4 mm pieces of EVA foam.
3. Glue the foam pads in the designed indents (inner part of the holder, in front of each threaded hole). This prevents damaging the aluminum sleeve of eVOLVER when fixing the tube holder.
4. Screw the headless bolts in the holder until they touch the foam pads (visible when the pads slightly deform).
5. Slide the assembled cap holder on an eVOLVER sleeve.
6. Place a vial inside the sleeve and fit the vial cap to the cap holder (Fig. 2).
7. When the vial is at the desired position, tighten the screws of the holder to fix it in place.

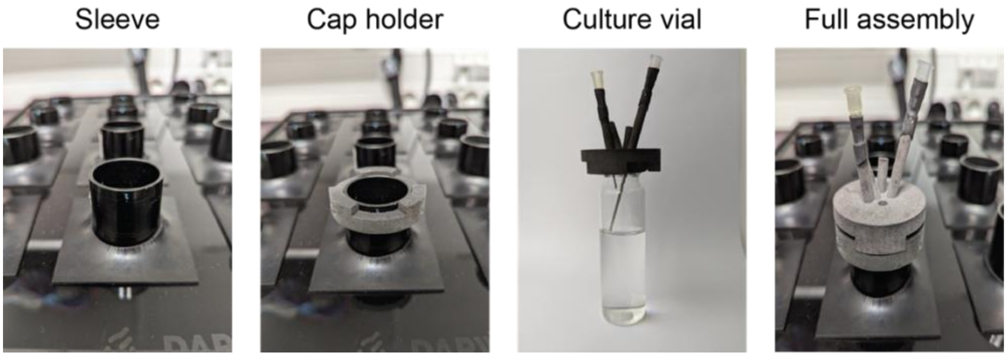

**Figure 2.** Assembly of the culture vials.

#### 4. STERILIZATION

Growing cultures in eVOLVER can be maintained for hundreds of generations. It is therefore necessary to take all possible measures for preventing contamination of the cultures. A crucial step in this process is the use of autoclaved bottles and vials.

---

**Media bottles**

1. Rinse the bottles and assembled bottle caps thoroughly with ultrapure water.
2. Run ultrapure water through both the tubing of the bottles and the caps.
3. Wash all elements using a standard laboratory glassware washer.
4. Let all parts dry completely (at room temperature or in an oven).
5. Check the proper connection of all bottle cap tubing (see Section 3) and close the bottles, leaving some play in the caps for the autoclave cycle. Cover the connections with aluminum foil.
6. Autoclave the bottles using a standard glass cycle.
7. Attention: to prevent contaminations, ensure that the sterilized tubing never comes in contact with any surface.
8. Following the sterilization cycle, growth medium can be directly filtered in the bottles using standard filtration devices. Pay particular attention to maintaining the sterility of the environment throughout the filtration procedure.
9. To connect a bottle to eVOLVER, tear the top part of the foil and spray with 70% ethanol, remove the luer lock cap and connect it rapidly to the appropriate pump line.

---

**Vials**

1. Rinse the vials thoroughly with ultrapure water.
2. Run ultrapure water through the cap needles.
3. Wash the vials using a standard laboratory glassware washer.
4. Let the vials dry completely (at room temperature or in an oven).
5. Put a magnetic stirring bar in each vial.
6. Assemble the vials and caps (see Section 3).
7. Cover the cap connections with aluminum foil.
8. Autoclave the vials using a glass cycle.
9. Keep the aluminum foil covering the connections until putting the vials into the eVOLVER sleeves to ensure sterility.

---

**Tubing**

Requirements:

- 2 L beakers
- 70% ethanol
- Sodium hypochlorite or bleach
- Ultrapure water
- Media bottles

Before starting any eVOLVER experiment, it is imperative that all the tubing that connects the different elements (*e.g.*, influx and efflux lines) is thoroughly cleaned. This step is critical for conducting experiments that can last for several months, limiting the potential sources of contamination in the cultures.

1. Wear a lab coat and gloves to protect yourself from the bleach.
2. If necessary, connect tubing splitters with the inputs of the influx pumps according to the specific setup of the experiment.
3. Prepare two 2 L beakers with 0.5 L of 1:10 bleach or 0.5 L of ultrapure water with 18 mL sodium hypochlorite (~14%) in each.
4. In the first beaker, immerse the inputs of the influx pumps. In the second beaker, immerse the outputs of the influx pumps and the inputs of the efflux pumps. Place the output tubing of the efflux pumps in a waste tank.
5. Start eVOLVER by turning on the 5V and then 12V switches (always switch the 5V first).
6. Start the pump array.
7. Using the eVOLVER touchscreen, click on *SETUP* and navigate to parameter *Peristaltic pump array*.
8. Click on the *IN1* (*i.e*. array of influx pumps) and *E* (*i.e*. array of efflux pumps) toggles and select all vials.
9. Use the slider to set a running time of 20 s.
10. Press on the button *Pump: 20s* to run the pumps. For optimal sterilization, make sure that no air goes inside the tubing.
11. Repeat this step two more times.
12. After pumping bleach for the last time, keep it inside the tubing for 30 min to overnight.
13. Run the pumps until both beakers are empty. Run again to remove bleach leftovers.
14. Prepare two 2 L beakers with 0.5 L of 70 % ethanol.
15. Place the tubing inside the ethanol beakers as in step 4.
16. Run the pumps three times for 20 s.
17. Connect the input tubing of the **influx** pumps to the media bottles. Keep the output tubing of the influx pump and the input tubing of the efflux pump in an empty beaker.

18. Run the **influx** and **efflux** pumps twice for 20 s to remove the ethanol. Following this procedure, the influx lines will be filled with medium and the efflux lines with air.
19. Spray ethanol on the Luer connectors and connect them to the needles of the vials.

### 5. CALIBRATIONS

#### 5.1. Temperature calibration

The temperature of each eVOLVER culture is controlled independently. This is achieved using a feedback control combining two electronic elements that heat the aluminum tube of the eVOLVER sleeve and a thermistor that monitors the temperature of this tube. The system must therefore be calibrated in order to determine the relation between the temperature of the aluminum tube and that of the culture. This section describes how to calibrate the temperature control of each eVOLVER sleeve.

---

**Before starting**

The calibration of the temperature control is accomplished by measuring three different temperatures using an external thermometer directly immersed in the medium, and associating the obtained values to the corresponding measurements acquired by the thermistor. This procedure is repeated for each individual eVOLVER sleeve. The approximate target temperatures are room temperature (RT), RT + 6 ºC and RT + 12 ºC. For this range, the provided eVOLVER application must be used. The following protocol includes ample waiting times to ensure temperature equilibration at each step. The user may choose to reduce these waiting times if equilibrium is reached faster.

Requirements:

- 16 eVOLVER glass vials
- eVOLVER computer
- High accuracy temperature probes

---

**Protocol**

1. Fill the 16 glass vials with 20 mL of water and place them in the eVOLVER sleeves.
2. Assemble eVOLVER with all the components used for a standard experiment, except for the media tubing. This will improve the accuracy of the calibration.
3. Start eVOLVER using the 5V and 12V switches (always switch the 5V first).
4. Start the eVOLVER electron app on the computer.
5. Click on *CALIBRATIONS* > *TEMP*.
6. Enter a name for this specific calibration and write it down. This name will be used later to generate the temperature calibration curves.
7. Click on *Start Temperature Calibration*. This will turn off the sleeve heaters.
8. Let eVOLVER equilibrate to RT for ~1 hour.
9. Introduce a temperature probe in the water through the sampling port and let the temperature measurement equilibrate.
10. Enter the temperature measured with the external probe in the eVOLVER electron app.
11. Repeat steps 9 and 10 for all vials.

*Note:* For each of the three target temperatures, eVOLVER records the thermistor measurements of all vials at the end of the process, once the values for the water have been entered for each vial. To limit temperature variations and increase the calibration accuracy, we suggest to use several temperature probes and simultaneously measure the water temperature for several vials. This will reduce the time between the external measurements and the acquisition of the thermistor readings.

12. In the eVOLVER app, click on the center button (RT). This will record the temperatures measured by the thermistors. During this process, the app displays *"Collecting raw values from eVOLVER…"*.
13. Click on the right button (+6 ºC) to move to the next calibration step.
14. Click on the center button (RT + 6 ºC) for eVOLVER to set the new temperature. Let the temperature equilibrate over the day.
15. Repeat steps 9 to 11.
16. In the eVOLVER app, click on the center button (RT + 6 ºC) for eVOLVER to record the temperatures measured by the thermistors.
17. Click on the right button (+6 ºC) to move to the next calibration step.
18. Click on the center button (RT + 12 ºC) for eVOLVER to set the new temperature. Let equilibrate overnight.
19. Repeat steps 9 to 11.
20. In the eVOLVER app, click on the center button (RT + 12 ºC) for eVOLVER to record the temperatures measured by the thermistors.
21. Click on the right button (pen icon) to save the calibration data.

---

**Calculating the calibration curves**

1. Open a Terminal and go to the DPU folder within the Terminal.
2. Use the script *calibrate.py* to generate the calibration curves.

```
# Calculate temperature calibration curves
python calibration/calibrate.py -a <ip_address> -n <file_name> -t linear
-f <name_after_fit> -p temp
```

Replace the following parameters:

- *<ip_address>*: IP of the eVOLVER machine.
- *<file_name>*: name of the calibration determined in step 6 of the calibration section above.
- *<name_after_fit>*: name of the generated calibration curves that will be used during an eVOLVER experiment.

3. If the displayed calibration curves are of good quality, type *"y"* and press *"Enter"* to update eVOLVER with the new temperature calibration. This specific calibration will be named after the parameter *name_after_fit* used in the command above.
4. To activate the calibration, use the eVOLVER touchscreen or the eVOLVER app: navigate to *SETUP* and select the calibration name next to the *TEMP* label.

---

**Improving the temperature control**

To fine-tune the temperature control, the user can check any potential remaining offset between the temperature indicated by eVOLVER after calibration and that determined using a probe directly immersed in the culture medium.

1. Start the eVOLVER using the 5V and then 12V switches.
2. On the eVOLVER touchscreen, click on *SETUP*.
3. Select the appropriate temperature calibration.
4. Select all vials and set them to the working temperature of the experiment.
5. Let the temperatures equilibrate in all vials.
6. Introduce a temperature probe through the sampling port of the first vial and let the temperature measurement equilibrate.
7. Write down the temperature of the probe and that displayed on the eVOLVER screen.
8. Repeat steps 6 and 7 for each vial.
9. For each sleeve, calculate the offset between the real temperature and that determined by eVOLVER.
10. When configuring the experimental parameters in the *custom_script.py* (see Section 6.2), correct the temperature settings of all vials using their respective offsets.

---

**Modifying the temperature calibration range**

If necessary, the temperature calibration range can be modified.

1. Open *eVOLVER_computer/evolver-electron/app/containers/TempCalibrate.js* in a code editor.
2. Edit line "*deltaTempRange: [0, 700]*" to modify the temperature range (testing may be required to determine the exact values that must be provided to the Arduino to obtain the correct range).
3. Recompile the app as follows :
   - Open a Terminal and navigate to the app folder: *cd eVOLVER_computer/evolver-electron.*
   - Run the command *yarn* to install dependencies.
   - Run the command *yarn package* to compile the new eVOLVER app.

The newly compiled app can be found in the *release* folder inside the *evolver-electron* folder. The compiling should be performed with the same operating system as the one used to run the app. More detailed instructions can be found at https://github.com/FYNCH-BIO/evolver-electron.

#### 5.2. Pump calibration

The eVOLVER system automatically dilutes the cultures using a set of 12V peristaltic pumps. When a dilution is needed, the eVOLVER DPU calculates the bolus volume and activates the pumps. The pumps are operated on a time-based strategy. Thus, the time required to inject the necessary volume of medium depends on a calibration that must be performed for each pump prior to the evolution assay.

---

**Before starting**

The calibration of the eVOLVER pump system is an estimation of the flow rate of each individual pump. This is achieved by measuring the volume pumped in a determined amount of time. Prior to calibrating the pumps, the following actions should be performed:

- Spray some glue (see Section 8.5) on the axes of the pump motors to improve adhesion with the rotating barrels. Let it dry prior to use.
- Activate the pumps for ~ 1 min to ensure that they all work.
- If the pumps have not been used for a long time, perform a tubing cleaning procedure (see Section 4).

Requirements:

- 16 plastic tubes (15 or 50 mL)
- 1 L beaker filled with water
- 10 mL plastic pipette and automatic pipette

---

**Protocol**

1. Fill a beaker with water, immerse the inputs of the influx pumps, and place their outputs in an empty beaker.
2. Start the eVOLVER using the 5V and then 12V switches.
3. Start the pump array.
4. Using the eVOLVER touchscreen or the eVOLVER app, click on *SETUP* and navigate to the *Peristaltic Pump Array* parameter.
5. Click on the *IN1* (*i.e.* array of influx pumps) toggle and select all vials.
6. Run the pumps for 2 min to fill the lines with water. Check that all pumps are turning.
7. Run the pumps again for 20 s to check that water is flowing through all outputs.
8. Place each output inside a separate 15 or 50 mL tube and run the pumps for 15 s.
9. Measure the volume of water in each collection tube using a graded pipette.

*Note*: all lines should pump around 10 mL in 15 s. Lines that pump volumes lower than 8.5 mL in 15 s should be checked.

10. Repeat the process three times.

11. If the volumes pumped in any of the repeats differ by more than 0.5 mL, repeat again, and discard the outlier.
12. Calculate the average flow rate (average pumped volume / 15 s).

---

**Applying the calibration**

Create a file called *pump_cal.txt* containing a single row with all the 16 calculated flow rates separated by tabs. Alternatively, export the flow rates as tab-separated text from any spreadsheet software. This file should be stored inside the experiment folder (see Section 2.3).

*Note:* the eVOLVER DPU does not need the flow rate of the efflux pumps, as they are programmed to always run for 20 s longer than the influx pump, preventing any overflow. Nevertheless, we advise the user to regularly repeat the calibration process for the efflux pumps to check that they are working properly and that the flow rates are similar to those of the influx pumps. Failure of the efflux pumps is one of the most common sources of flooding.

##### 5.3. OD calibration

eVOLVER evaluates the optical density (OD) of each culture by measuring the light scattering of an infrared (IR) LED using photodiodes placed at angles of 90º and 135º. A calibration is thus needed to establish, for each eVOLVER sleeve, the relation between scattered light and OD. This is achieved by associating the raw scattering values measured by eVOLVER with ODs provided by an accurate benchtop spectrophotometer. For a working $OD_{595}$ range between 0.2 and 0.5 (standard range for exponentially growing yeast cultures), only the 135º photodiode is used. This section describes how to calibrate the optical density measurements of the eVOLVER sleeves.

*Note:* for simplicity, although eVOLVER does not directly measure optical density, we refer to these measurements as "OD". "$OD_{595}$" is only used in this guide when direct measurements are performed using a spectrophotometer.

Requirements:

- 16 clean eVOLVER glass vials
- 1 L pre-warmed Phosphate Buffered Saline (PBS)
- 10 mL plastic pipette and automaticpipette
- 1 mL pipette tips
- Long syringes or vacuum pump

---

**eVOLVER preparation**

1. Start the eVOLVER using the 5V and then 12V switches.
2. If needed, check the section *Optimizing LED power* below to set the appropriate LED power for your eVOLVER system.
3. Using the eVOLVER touchscreen, click on *SETUP* and select all the vials.
4. Select the appropriate temperature calibration (see Section 5.1).
5. Navigate to the stir option interface and set it to 10.
6. Navigate to the temperature option interface and set it to the appropriate value.
7. Clean the outside of fully assembled autoclaved vials (see Section 4) with 100% isopropanol and a dust-free cloth.
8. Fix the vials in the sleeves using the new cap system (see Section 3).

*If the calibration is followed by an experiment*:

- Make sure to keep sterile conditions throughout the whole process. Use new sterile tips for every OD point or, if using a repeater pipette, flame the tip before each inoculation.
- Go through the full tubing cleaning procedure, including the last step of washing out the ethanol with the experiment medium (see Section 4).
- Connect the influx and efflux tubing once they are clean, and ensure that they are properly tightened.
- Place the output of the efflux tubing in a liquid waste container with bleach.

9. Fill all vials with 20 mL of pre-warm filtered PBS using a pipette fitted with a 1mL tip.
10. Ensure that all the stirring bars are rotating properly:
    - Using the eVOLVER touchscreen, click on *SETUP* and select all the vials.
    - Navigate to the stir option interface and set it to 0 to stop stirring.
    - Set it back to 10 to resume stirring.

---

**Cell preparation**

The amount of cells used for OD calibration is calculated based on the working OD range of the experiment. When the calibration is followed by an experiment, the cells used for the calibration in each vial should be the same as those that will be used for the experiment. Separate cultures of distinct strains may thus be necessary. This protocol is optimized to calibrate eVOLVER using the 135° photodiode, around a working $OD_{595}$ range between 0.2 and 0.5.

1. Grow 500 mL of cells to an $OD_{595}$ of ~0.45.
2. Spin the cells for 5 min at 3000 rpm and wash them once with 50 mL of pre-warmed filtered PBS.
3. Spin the cells again and concentrate them to a final volume of 20 mL in PBS (around 10 OD).
4. Measure the $OD_{595}$ of the concentrated culture with a benchtop spectrophotometer (dilutions should be used to be in the linear range of the spectrophotometer and obtain a more accurate determination of the density).
5. Use the provided Excel file (*Cal_Inoculation.xlsx*) to get the 16 OD points for the OD calibration.
6. If necessary, modulate the OD range by modifying the volumes inoculated at each step in the Excel file.

*Note*:

- Increasing the density of OD points near the zero as well as in the working range of the experiment will improve the quality of the calibration.
- At this point, the ODs provided in the top table of the Excel file are only estimations based on the OD of the stock culture and the theoretical dilutions at each inoculation step. These ODs will be adjusted at the end of the calibration procedure (see below).

---

**Calibration**

1. Start the eVOLVER electron app on the computer.
2. Click on *CALIBRATIONS* > *OD*.
3. Enter a calibration name and write it down. This name will be used later to generate the OD calibration curves.
4. Enter the 16 ODs calculated using the provided Excel file, in ascending order.
5. Click on the center button (Play icon) to start recording the blank OD point (PBS).
6. Wait until the center button displays a checkmark.
7. Click on the right arrow to move to the next OD point.

8. Inoculate cells to the 16 vials according to the first step of the calibration (see *Cal_Inoculation.xlsx* Excel file) and wait for 20 s.
9. Click on the center button (Play icon) to record the new OD point.
10. Click on the right arrow to move to the next OD point.
11. Repeat steps 8 to 10 for the 16 OD points.
12. Click on the right button (Pen icon) to save the calibration data.

---

**Correcting the calibration data**

For accurate calibration, the final $OD_{595}$ of each vial must be measured to correct the theoretical values calculated in the Excel file.

1. Sample 1 mL from each vial and measure the $OD_{595}$ using a benchtop spectrophotometer (dilute if necessary to be in the linear range of the machine).
2. Enter these final ODs in the second table of the Excel file. These values are automatically averaged and the initial $OD_{595}$ of the stock culture is back-calculated using the dilution factor. The bottom "*OD back calculation*" table now provides a new set of back-calculated ODs for each inoculation step.
3. On the computer, create a new folder with the name of your calibration. This folder will be used to save a copy of the *calibrations.json* file and to correct the calibration data.
4. Open a Terminal and navigate to this new folder within the Terminal.
5. Run the following command to download the *calibrations.json* file. It will ask for the password of the eVOLVER server provided by the manufacturer.

```
# Download calibrations.json from the device using scp
scp pi@<ip_address>:evolver/evolver/calibrations.json calibrations.json
```

Replace the parameter *<ip_address>* with the IP of the eVOLVER machine.

6. Open a Terminal and go to the *evolver-utils* folder (see Section 2.2) within the Terminal.
7. Run the *Cal_correction.py* script, providing the 16 back-calculated ODs of each inoculation step (see below). This will output a *calibrations.json.out* file with the modified calibration data in the folder created in step 3.

```
# Correct the calibration data with the back-calculated ODs
python Cal_correction.py -c <path_to_calibrations_json> -n
<calibration_name> -l <OD_0> <OD_1> ... <OD_15>
```

Replace the following parameters:

- *<path_to_calibrations_json>*: path to the file downloaded in step 4.
- *<calibration_name>*: name of the calibration determined in step 3 of the calibration section above.
- *<OD_0>, <OD_1> ... <OD_15>*: 16 OD values back-calculated in the Excel file separated by a single space.

8. Open a Terminal and go to the folder created in step 3 within the Terminal.
9. Upload the new *calibrations.json.out* file to eVOLVER, overwriting the old *calibrations.json* file as follows:

```
# Upload calibrations.json.out to the device using scp
scp ./calibrations.json.out
pi@<ip_address>:evolver/evolver/calibrations.json
```

Replace the parameter *<ip_address>* with the IP of the eVOLVER machine.

---

**<u>Generating the OD calibration curves</u>**

1. Open a Terminal and go to the DPU folder within the Terminal.
2. Run *calibrate.py* to calculate and display the calibration curves. A *pyplot* window will appear.

```
# Calculate OD calibration curves
python calibration/calibrate.py -a <ip_address> -n <file_name> -t sigmoid
-f <name_after_fit> -p od_135
```

Replace the following parameters:

- *<ip_address>*: IP of the eVOLVER machine.
- *<file_name>*: name of the calibration determined in step 3 of the calibration section above.
- *<name_after_fit>*: name of the calibration curves that are generated and that will be used during the eVOLVER experiment.

3. The calibration curves can be saved as a pdf or a png file in the calibration folder for future reference. Click on the Save button (floppy disk icon) and type the name of the calibration with the appropriate file extension.

4. If the displayed calibration curves are of good quality, type *"y"* and press *"Enter"* to update the eVOLVER with the new calibration. This calibration will be named after the parameter *name_after_fit* used in the command above.

> *Note*: Make sure there is medium flowing out of the vials through the efflux tubing. This validates that the medium is at the maximum level, as determined by the height of the efflux needles, inside the vials.

5. To activate the calibration, navigate to *SETUP* in the eVOLVER touchscreen or the eVOLVER app and select the calibration name next to the OD label.
6. If the calibration is not followed by an experiment, turn off the 12V and then 5V switches, remove the vials and clean all the parts.

---

**Starting an experiment right after a calibration**

To optimize an eVOLVER experiment, we recommend performing a new OD calibration immediately prior to each evolution assay using, for each vial, the cells that will also be used for the experiment.

1. Once the OD calibration procedure is complete, turn off the stirring via the touchscreen and remove the cultures from the vials using long syringes or a vacuum pump.
2. Using the eVOLVER touchscreen or the eVOLVER app, click on *SETUP* and navigate to the *Peristaltic Pump Array* parameter.
3. Click on the *IN1* (*i.e*. array of influx pumps) and *E* (*i.e*. array of efflux pumps) toggles and select all vials.
4. Use the slider to select 20 s and press the button *Pump: 20s*. Repeat this three times.
5. Turn on the stirring: navigate to the *Stir Rate* parameter, use the slider to select a value of 10, and press the button *Stir: 10*.
6. Wait for 5 minutes.
7. Turn off the stirring: in the *Stir Rate* parameter, use the slider to select a value of 0, and press the button *Stir: 0*.
8. Remove all liquid from the vials with long syringes or a vacuum pump.
9. Wash out any residual cell from the calibration by repeating steps 2 to 8.
10. Fill the vials with the medium that will be used for the experiment by repeating steps 2 to 4.
11. Continue with the standard experimental procedure (see Section 6).

---

#### Optimizing LED power

To improve the dynamic range of the OD calibration curves, it is recommended to perform several OD calibrations using different LED power values to find the optimal one for each individual sleeve (see main manuscript). This section details how to modify the LED power of each sleeve and explains how to determine the optimal values.

> *Note*: although the general principles explained here are universal for different configurations of the eVOLVER framework, the values provided are appropriate for optimizing the LED power of a system with a 51 MOhm resistor pack, using the 135º photodiode for the light scattering recording.

1. Start the eVOLVER using the 5V and then 12V switches.
2. Use the script *Cal_LEDpower.py* to set the LED powers. As a starting default, use 2060.

```
# Set the LED power
python Cal_LEDpower.py -a <ip_address> -p <power>
```

Replace the following parameters:

- *<ip_address>*: IP of the eVOLVER machine.
- *<power>*: power for all the 16 LEDs. Use 2060 as default.

3. Restart the eVOLVER to apply the changes.
4. Perform an OD calibration following the sections above, from *eVOLVER preparation* to *Generating the OD calibration curves*. In this last section, save the calibration plots (step 3) but do not update the calibration to eVOLVER (in step 4 type '*n*' and press "*Enter*").
5. Repeat the LED optimization steps two more times, using powers 2062 and 2064.
6. Compare all three calibration plots and, for each sleeve, identify which power produces curves with a better dynamic range in the appropriate OD working range.
7. Run the script *Cal_LEDpower.py* to set the final LED power for each individual sleeve.

```
# Set the LED power
python Cal_LEDpower.py -a <ip_address> -p <power0> <power1> … <power15>
```

Replace the following parameters:

- *<ip_address>*: IP of the eVOLVER machine.
- *<power0>, <power1> ... <power15>*: LED power of each sleeve, separated by a single space.

8. Restart the eVOLVER for to apply the changes.
9. Repeat with alternative power values if the results above are not satisfactory.

**Deleting old calibrations**

Since it is advised to perform an OD calibration before each experiment, the eVOLVER system will accumulate obsolete calibrations. This section describes how to delete old temperature and OD calibrations using the provided script.

1. On the eVOLVER computer, create a new folder. This folder will be used to save a copy of the *calibrations.json* file and to delete the old calibrations.
2. Open a Terminal and navigate to this new folder within the Terminal.
3. Run the following command to download the *calibrations.json* file. It will ask for the password of the eVOLVER server provided by the manufacturer.

```
# Download calibrations.json from the device using scp
scp pi@<ip_address>:evolver/evolver/calibrations.json calibrations.json
```

Replace the parameter <*ip_address*> with the IP of the eVOLVER machine.

4. Open a Terminal and go to the *evolver-utils* folder (see Section 2.2) within the Terminal.
5. Run the script *Cal_delete.py* with the -*l* parameter to display a list of available calibrations.

```
# List all available calibrations
python Cal_delete.py -l
```

6. Write down the name of the calibrations to be deleted.
7. Run the script *Cal_delete.py* with the -*n* parameter to delete specific calibrations.

```
# List all available calibrations
python Cal_delete.py -n <cal_name>
```

Replace the parameter <*cal_name*> with the name of the calibration to be deleted.

8. The Terminal will prompt *Delete? (y/n)*. To confirm, type *"y"*. This will overwrite the *calibrations.json* file downloaded in step 3.

```
# Upload calibrations.json to the device using scp
scp ./calibrations.json pi@<ip_address>:evolver/evolver/calibrations.json
```

Replace the parameter <*ip_address*> with the IP of the eVOLVER machine.

9. Repeat the procedure for all calibrations that need to be deleted.

### 6. RUNNING AN EXPERIMENT

eVOLVER is a powerful system to monitor and control continuous cultures of microbial populations. Running an eVOLVER experiment requires thorough planning and a multi-step procedure, from the calibration of the framework to the modification of the experimental scripts. In this section, we provide a step-by-step protocol to start and monitor an eVOLVER experiment.

**Checklist before starting an experiment**

- According to the number of sleeves (*n*) that will be used, assemble and sterilize at least *n*+2 complete vials (see Sections 3 and 4).
- The autoclaving process sometimes alters the original shape of the vial caps, preventing the proper fitting of the vial in the sleeve. Verify this beforehand and adjust if necessary.
- Prepare all the necessary bottles with medium. Always make sure that all the tubing is properly connected.
- Prepare splitters according to the experimental setup.
- Ensure that all the tubing is clear and unclogged, and that the pumps are working.

---

#### 6.1. eVOLVER startup

The first steps of this protocol are identical to those described in the OD calibration procedure, as we encourage the user to perform a calibration prior to each experiment.

1. Start the eVOLVER using the 5V and then 12V switches.
2. Using the eVOLVER touchscreen, click on *SETUP* and select all the vials.
3. Select the appropriate temperature calibration.
4. To set the stirring, navigate to the *Stir Rate* parameter, select a value of 10, and press the button *Stir: 10.*
5. Navigate to the *Temperature* parameter, set it to the appropriate value and press the *Temperature* button.
6. Clean the outside of the fully assembled autoclaved vials (see Section 4) with 100% isopropanol and a dust-free cloth.
7. Fix the vials in the sleeves using the new cap system (see Section 3).
8. Start the OD calibration procedure (see Section 5.3) or select the appropriate OD calibration in the eVOLVER app.

*Note*: if an old OD calibration is used, the user must adjust the vials so that the raw values displayed in the eVOLVER screen are as close as possible to the calibration raw zero values in the *od_raw_zero.json* file.

##### 6.2. Modifying the experiment code

An eVOLVER experiment is controlled by two scripts inside the experiment folder:

- *eVOLVER.py* has all the logic to start, pause and stop an experiment, as well as to communicate with the eVOLVER machine.
- *custom_script.py* contains the main settings and the logic of the turbidostat experiment designed by the user.

Before a new experiment:

1. Duplicate the template folder and change its name to an experiment-specific name (see Section 2.3). The *custom_script.py* then needs to be modified, using standard free editors such as Notepad, Sublime Text or VS Code, as follows.
2. Change the experiment name *EXPT_NAME (line 14)*. This name must end with "*_expt*" to be recognized by the graphing tool.
3. Select the eVOLVER machine that will be used by specifying its IP address *EVOLVER_IP (line 15).*
4. Set the temperature of the vials *TEMP_INITIAL (line 20)*

*Note:* to modify the temperature parameter during an experiment, changing the values of *TEMP_INITIAL* will not work, as this parameter only sets the initial temperatures of the vials. This can however be achieved by manually modifying the file *vialX_temp.txt* (where X represents the vial number) inside the *temp* folder (see Section 2.3).

5. Set the stirring speed *STIR_INITIAL (line 24).*
6. Set the volume of the vials *VOLUME (line 28).*
7. Set the name of the pump calibration file *PUMP_CAL_FILE (line 29).*
8. Check that *OPERATION_MODE = "turbidostat"* (line 30).
9. Edit the turbidostat parameters within the *turbidostat* function *(line 35)*: set the low and high OD thresholds (*lower_thresh* and *upper_thresh; lines 44 and 45).*
10. Set the *time_out* (additional time during which the efflux pumps will run to avoid overflow, *line 58*) and the *pump_wait* (time to determine whether a dilution was successful before triggering an additional round of dilution, *line 59*). We have reliably used 20 s and 3 min for the *time_out* and *pump_wait*, respectively.

*Note:* All these scripts are provided with the default experimental parameters that were found to be optimal for evolving exponentially growing fission yeast cells. They should be modified according to the experimental design.

##### 6.3. Starting the experiment

1. Make sure the Telegram alert system is configured (see Section 8.4).
2. If using the scale system (see Section 6.4), tare each scale by placing an empty bottle of the same capacity and press the scale button for 5 seconds until the blue light on the button stops blinking. Subsequently place each media bottle on its respective scale.
3. Open a Terminal and go to the experiment folder within the Terminal.
4. Run the following command to start the experiment.

```
# Start the experiment script
python eVOLVER.py
```

5. The Terminal will prompt *"Calibrate vials to blank? (y/n)"*. Type *"y"* and press *"Enter"*.
6. The Terminal will prompt *"Use raw blank instead of OD blank? (y/n)"*. Type *"y"* and press *"Enter"* to blank the vials using a raw blank (see main manuscript).
7. Open a new Terminal and go to the DPU folder within the Terminal.
8. Run the following command to start the graphing tool.

```
# Start the graphing tool
python graphing/src/manage runserver
```

9. On a web browser in the eVOLVER computer, navigate to the address http://127.0.0.1:8000 to display the graphing tool.
10. Wait until the OD readings are stable at OD=0 and the temperature readings are stable at the set values.
11. Stop the experiment by pressing "*Ctrl + C*" twice within the experiment Terminal (step 3) and delete the *Experiment_expt* folder.
12. Start a new experiment by repeating steps 4 to 6.
13. Pause the experiment by pressing *"Ctrl + C"* within the experiment Terminal (step 3).
14. Concentrate cells to 1 mL and inoculate them through the sampling port. The amount of cells to inoculate should be calculated based on the starting $OD_{595}$ that is targeted by the user.
15. In the experiment Terminal, press *"Enter"* to resume the experiment.
16. Open the graphing tool on a web browser and check that the cultures show the expected OD.

##### 6.4. Monitoring an experiment

eVOLVER takes advantage of a graphical interface to display the experimental data. This interface can be accessed through a web browser on the eVOLVER computer (see above). Here we describe how to start, configure and take advantage of this graphical interface to monitor an eVOLVER experiment.

**Configuring the scale interface**

We take advantage of a scale system that monitors the weight of eVOLVER media bottles (see below). A minor configuration is needed to integrate the data from the scale system in the graphing tool:

1. Open the file *scales_config.json* in the experiment folder.

```
{
    "probes": [1111, 2222, 3333],
    "conditions": {
    1111: "Condition 1",
    2222: "Condition 2",
    3333: "Condition 3",
    }
}
```

*scales_config.json*

2. Modify the parameter *probes* with a list of IDs for the purchased scales.
3. Modify the parameter *conditions* with an array of *keys: values*, where the keys are the scale IDs, and the values are the name of the scale or media bottle between double quotes.
4. Open the *creds.json* file in the experiment folder.
5. Modify the parameter *bker_user* with the username provided by the scale manufacturer.
6. Modify the parameter *bker_pass* with the password provided by the scale manufacturer.

---

**Modifying the graphing parameters**

Other parameters of the graphing tool can be modified, such as the axis range of some graphs and the size of the sliding window used to calculate population generation times. We have selected the most useful parameters and provide further instructions within the graphing tool code.

1. Open the file *graphing/src/cloudevolution/views.py* located in the DPU folder.
2. Navigate to the comments starting with *# NOTE* and follow the instructions.

---

**Starting the monitoring interface**

1. Open a Terminal and navigate to the DPU folder of the experiment within the Terminal.
2. Run the following command to start the graphing tool:

```
# Start the graphing tool
python graphing/src/manage runserver
```

3. This command activates the monitoring interface, which can then be accessed by navigating to the address https://127.0.0.1:8000 in any web browser installed on the eVOLVER computer.

---

#### Using the monitoring interface

##### Experiment page

Clicking on the experiment name on the sidebar opens the experiment page, where a summary of the experiment is displayed, including the generation time graphs for all the vials of the eVOLVER.

##### Vial page

To access individual vial pages, click on the vial number button in the top navigation bar. The page will display (1) an OD graph, (2) a generation time graph, and (3) a temperature graph for the selected vial. Moreover, footnotes provide the last time data have been acquired.

##### Dilutions page

This page displays information about (1) the total amount of medium consumed by each vial since the start of the experiment (as calculated from the total number of dilution events), (2) the pump efficiency (% efficiency = (number of dilution intervals – number of additional dilution steps applied) / number of dilution intervals), and (3) the amount of medium consumed from each bottle that is currently connected to eVOLVER (as calculated from the dilution events of all vials connected to that bottle).

Moreover, this page allows the user to define a bottle setup that associates each bottle to a set of vials and allows for the tracking of the bottle changes and media consumption:

1. Click on the button *Edit bottle setup*. A pop-up will appear, displaying a default *bottle0*.
2. Add the vial numbers separated by commas and the volume of the bottle in liters.
3. Click on the + button to create a new bottle and repeat step 2.
4. Click on the button *Save changes*. This will create a file *bottles.txt* inside the experiment folder.
5. To change a bottle during an experiment, refer to Section 7.2.

##### Scales page

We take advantage of a scale system that monitors the weight of eVOLVER media bottles, calculates the live consumption rates and alerts the user when the detected weight is lower than a predefined threshold (see main manuscript). The *Scales* page, which can be accessed for each experiment from the top navigation bar, displays: (1) the last weight measured by each of the scales configured in the *scales_config.json* file (see above) and (2) a set of graphs showing all the data measured by each scale for the duration of the experiment. If the latter is not displayed or to update the graphs, click on the button *GetScale*.

### 7. CONDUCTING AN EVOLVER EXPERIMENT

#### 7.1. Sampling during an experiment

Regularly sampling the eVOLVER cultures allows for monitoring the phenotype of the experimental populations and detect potential contaminations. Moreover, regular freezing of population aliquots is essential: it will serve as a “population snapshot” that may be used both to restart a failed culture (see below) or to perform further analyses after the experiment. Thus, the optimal sampling frequency will depend on the specifics of the evolution assay.

---

**Before starting**

The caps of the eVOLVER vials incorporate a sampling port of reduced diameter to limit contaminations. For this reason, to sample the cultures, we use 1 mL sterilized plastic syringes with 7 cm, 20G sterilized needles. To avoid contaminations, all measures should be taken to maintain a sterile environment (*e.g.* wear a lab coat and spray gloves with 70% ethanol before touching the vials).

---

**Protocol**

1. Take an empty 1 mL micro-pipette tip box and place the sampling needles in the box.
2. Autoclave the box using a standard glassware program.
3. Find the experiment Terminal on the computer and pause the experiment by pressing *“Ctrl + C”*.
4. Take a sterile syringe and attach it to one of the sampling needles.
5. Carefully insert the needle through the sampling port ensuring that the needle does not touch the cap.
6. Sample ~1 mL of culture into a 1.5 mL tube.
7. Once the sampling of all the cultures is done, press *“Enter”* in the experiment Terminal to resume the experiment.
8. Freeze the sample in the appropriate freezing medium.

#### 7.2. Changing a medium bottle

In eVOLVER, continuously growing cultures can be maintained for long periods of time, requiring periodic changes of the media bottles. This is an extremely important step during an eVOLVER experiment. This section describes the bottle changing procedure as well as the use of the system we have established to monitor all the bottle changes throughout the experiment.

---

**<u>Before starting</u>**

To avoid contaminations, keep the environment as sterile as possible, wear a lab coat, and spray gloves with 70% ethanol before touching the bottle connectors. Check that the new media bottles were sterilized properly and that the aluminum foil covers all the tubing.

---

**<u>Protocol</u>**

1. Find the experiment Terminal on the computer and pause by pressing "*Ctrl + C*".
2. If using the scale system, tare the scale with an empty bottle of the same capacity and then place the new media bottle on the scale.
3. Remove the aluminum foil from the new media bottle.
4. Spray 70% ethanol on the luer cap and connector of the new bottle as well as on the ones between the input of the influx pump and the old media bottle.
5. Place both bottles as close as possible to limit the exposure of the tubing to the environment when switching bottles.
6. Loosen the luer cap on both bottles and connect the input of the influx tubing to the new bottle. Make sure that the bottle is tightly connected.
7. Go to the "*Dilutions*" page on the web interface.

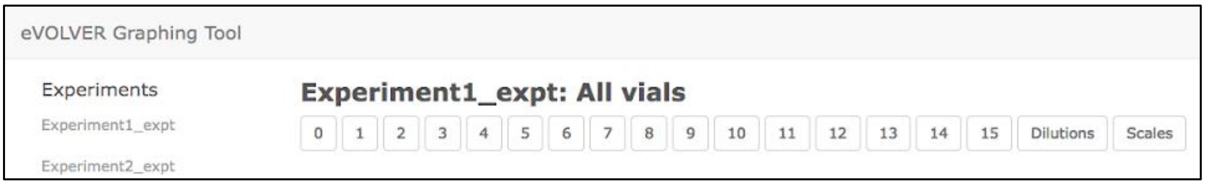

8. Click on "*See bottle setup*".

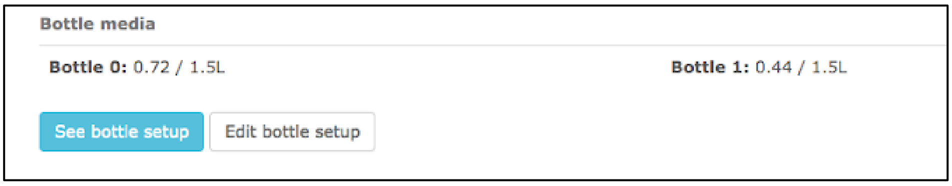

9. The current bottle setup will be displayed.

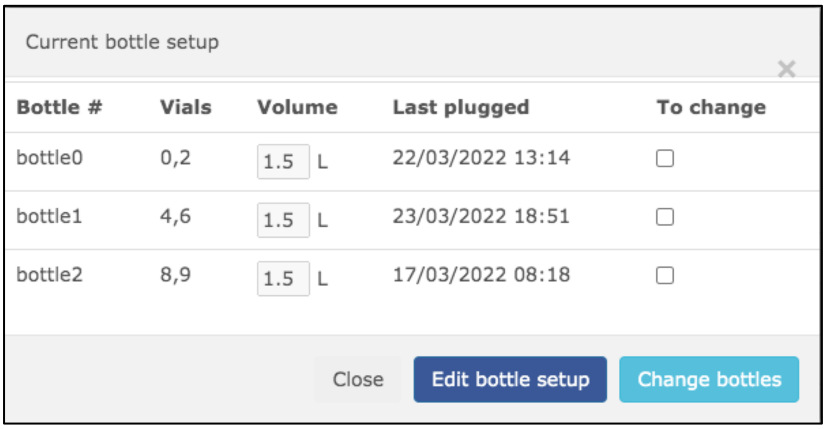

10. Click on the checkbox "*To change*" for the appropriate bottles.

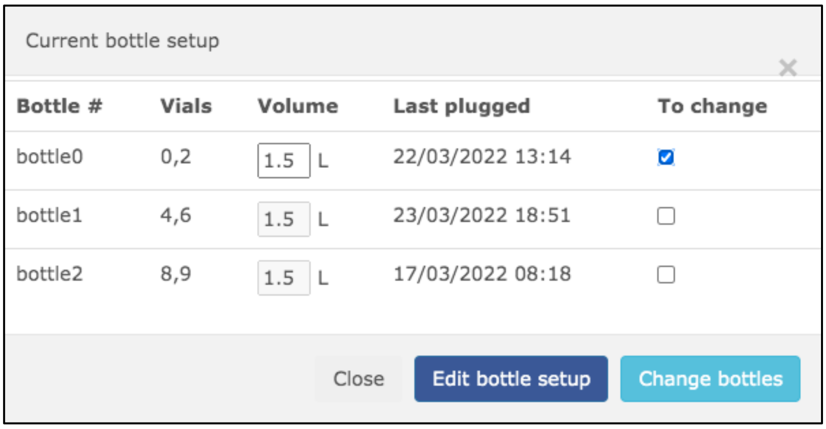

11. Modify the volume of the bottle if necessary and then click on *"Change bottles"*. This will log the time and the bottle information in the *bottles.txt* file inside the experiment folder (see Section 2.3).
12. In the experiment Terminal, press *"Enter"* to resume the experiment.

##### 7.3. Replacing problematic vials

Although eVOLVER is a robust culture system, hardware failures such as malfunctioning pumps due to normal wear, or clogging of the efflux tubing due to biofilm formation may occasionally happen. This can potentially cause a vial overflow that can leak onto the machine. Moreover, long-term growing cultures can become contaminated, in which case the corresponding vial should be replaced by a new vial containing a culture grown from the preceding frozen sample. Vial replacement is not straightforward as change in the glass vial itself alters the OD readings and interfere with the experiment (see main manuscript). Below we provide a guide and a set of tools to stop, remove and replace a problematic vial with limited impact on the overall ongoing assay.

---

**Removing a problematic vial**

1. Stop the main experiment completely by selecting the Terminal window and pressing *"Ctrl + C"* twice.
2. Open the *custom_script.py* and change the OD thresholds of the problematic vial to 9999. This prevents the automatic trigger of a dilution for this specific vial.
3. Resume the main experiment by running the following command:

```
# Resume the experiment
python eVOLVER.py
```

4. The system asks whether to continue the ongoing experiment. Type "*y*" and press "*Enter*" to continue. This allows to keep the experiment going while dealing with the failed culture.
5. Disconnect the problematic vial from the tubing and remove it from the sleeve.
6. Check the associated pumps, bottle and tubing. Clean if necessary (see Section 4).

---

**Restarting a problematic vial**

1. Grow cells from the latest frozen sample of the problematic culture.
2. Clean tubing as detailed in Section 4.
3. Place a new vial in the sleeve and connect it to the tubing.
4. Fill the vial with medium using the fluidic commands on the touchscreen or manually through the sampling hole. The use of pre-warmed medium is recommended.
5. Monitor vial temperature using the graphing tool until it equilibrates.
6. Copy the scripts *VialReplace_Equilibrate.py* and *VialReplace_Blank.py* from the *evolver_utils* folder into the experiment folder (next to *custom_script.py*).
7. Open a Terminal and navigate to the experiment folder within the Terminal.
8. Run the script *VialReplace_Equilibrate.py*.

```
# Display live OD and save it once equilibrate it
python VialReplace_Equilibrate.py <vial_number>
```

Replace the *<vial_number>* parameter with the number of the vial to be replaced. This script will:

- Display the calculated OD and raw values for the vial every 20 s.
- Save the equilibrated OD value for the vial in the file *equilibrated.json.*

9. Slightly rotate the vial and wait for the next measurement. Repeat this until the OD displayed is lower than 0.007. This step facilitates the calibration correction by matching the initial experimental blank with the blank of the new vial.
10. Pause the *VialReplace_Equilibrate* script using *"Ctrl + C"*. Follow the instructions displayed.
11. Repeat the steps 3 to 10 for all the vials that need to be restarted.
12. Run the script *VialReplace_Blank.py*.

```
# Apply new blank to experiment
python VialReplace_Blank.py
```

This script will:

- Make a backup of the *experiment.pickle* file containing the previous OD blank.
- Use the OD value after it is equilibrated (saved in the file *equilibrated.json*) to replace the original blank of the experiment.

13. Stop the main experiment completely by selecting the Terminal window and pressing *"Ctrl + C"* twice.
14. Open the *custom_script.py* and edit the OD thresholds of the replaced vial back to the target values.
15. Resume the main experiment by running the following command:

```
# Resume the experiment
python eVOLVER.py
```

16. The system asks whether to continue the ongoing experiment. Type *y* and press *"Enter"*.
17. Wait for a few measurements and check that OD = 0 using the graphing tool.
18. On the experiment Terminal, pause the main experiment by pressing *"Ctrl + C"*.
19. Inoculate the cells.
20. Press *"Enter"* to resume the main experiment.

### 8. APPENDIX

#### 8.1. Troubleshooting

**The eVOLVER screen turns white**

1. Stop the experiment by pressing *"Ctrl + C"* twice on the experiment Terminal.
2. Turn off the pump array, the 12V, and then the 5V switches of eVOLVER. Wait for a few seconds.
3. Restart the eVOLVER by turning on the 5V and then the 12V switches. Wait for the machine to complete its initialization process.
4. Start the pump array.
5. On the experiment Terminal, resume the experiment by running the following command:

```
# Resume the experiment
python eVOLVER.py
```

6. Continue from the existing experiment: when prompted with *"Do you want to continue from the existing experiment?"* type *"y"* and press *"Enter"*.
7. *Note*: Failing to follow the proper power-up sequence or not waiting for the initialization to restart the script may trigger extra dilutions, affecting the outcome of the experiment.

---

**The OD curve of a specific vial does not reach the desired lower threshold**

Some connections in the fluidic network may be loose. Check that the following parts are properly connected:

1. Inside tubing of the media bottle
2. Outside luer locks and tubing of the media bottle
3. Influx needle luer lock
4. Efflux needle luer lock

Alternatively, the influx or the efflux pumps may not be working properly. In this case, follow this procedure:

1. On the experiment Terminal, press *"Ctrl + C"* to pause the experiment.
2. Detach the pump head from the pump body. Clean both the internal parts and the rotating axis using 70% ethanol.
3. Spray some glue (see Section 8.5) on the axis of the pump motor to improve adhesion with the rotating barrels.
4. Connect the pump head again.
5. Resume the experiment by pressing *"Enter"* on the experiment Terminal.

If the problem persists, there may be a more significant problem with the fluidic system. The following can be tried:

1. Replace the output of the efflux tubing, as it may be clogged.
2. Replace the full pump head and tubing set with a new sterile part.

If the problem persists, there may be an issue with the calculation of the dilutions, likely due to wrong OD readings (*e.g.* change in vial positioning, biofilm formation). In this case, we suggest the following:

1. On the experiment Terminal, press *"Ctrl + C"* twice to stop the experiment.
2. Remove the vial from the sleeve for visual inspection.
3. Follow Section 7.3 to replace with a new vial if necessary.

---

**The OD readings are noisy**

The stirring bars may be jumping inside the affected vials, interfering with the OD readings.

1. Using the eVOLVER touchscreen, navigate to *SETUP*, select the affected vials, navigate to *Stir* and set it to 0.
2. Wait for a few seconds and similarly set the stirring back to the experiment value.

If there is no problem with the stirring, there may be a bacterial contamination:

1. Refer to Section 7.1 for sampling the affected culture.
2. Check the sample for any potential contamination using a microscope.

If there is no contamination, there may be a problem with the sleeve components:

1. Check the connection of the sleeve to the eVOLVER body.
2. Check the temperature readings, as this may also be noisy, suggesting a more general problem with the sleeve.
3. Check for any sign of medium overflow that could have damaged the electronic components.
4. Consider stopping the experiment if the problem is due to a significant overflow, as this could damage the eVOLVER.

#### 8.2. eVOLVER utils

Together with the main software to calibrate the eVOLVER system and run experiments, we provide a series of complementary tools to improve calibration and deal with vial failures. A number of these scripts have been mentioned previously. Here is a detailed explanation of the contents of the *evolver-utils* folder:

---

***Cal_inoculation.xlsx***

This Excel sheet allows for planning an OD calibration by (1) calculating the $OD_{595}$ range of the calibration, (2) back-calculating the $OD_{595}$ of the concentrated culture after measuring the ODs of each vial at the end of the calibration, and (3) back-calculating the $OD_{595}$ at each calibration step, taking into account the corrected $OD_{595}$ of the concentrated culture. Using this last set of ODs, the calibration can be corrected using the *Cal_correction.py* script.

---

***Cal_correction.py***

This script reorganizes the data of an OD calibration when running the new calibration protocol while using the original calibration software (fixed vials with successive inoculations of cells *vs* rotating vials of 16 different ODs). In addition, it updates the $OD_{595}$ values with the list of back-calculated ODs provided by the Excel sheet *Cal_Inoculation.xlsx*.

---

***Cal_LEDpower.py***

This script connects to eVOLVER and changes the pre-defined power of the IR LEDs used for cell density determination (see Section 5.3).

---

***Cal_delete.py***

This script allows the user to delete old calibrations from the eVOLVER *calibrations.json* file (see Section 5.3).

---

***VialReplace_Equilibrate.py***

This script facilitates the process of replacing a vial by displaying the live OD values obtained by eVOLVER while the user rotates the newly placed blank vial to achieve a position where the measured OD is close to zero. Once equilibrated, this script saves the current light scattering value in the *equilibrated.json* file, which will later be used to update the experiment blank with the script *VialReplace_Blank.py*.

---

***VialReplace_Blank.py***

After replacing a vial, allowing the OD equilibrate near zero, and using the script *VialReplace_Equilibrate.py*, this script uses the light scattering value stored in the *equilibrated.json* file to update the experiment blank of the selected vial. This allows for the compensation of any remaining differences in the OD measurement.

#### 8.3. eVOLVER hardware

We developed several hardware elements that protect the machine's electronic components from potential medium overflows, avoid unintended switching off the eVOLVER, and improve the reliability of the cell density measurements. Here we detail the different hardware parts and how to integrate them within the eVOLVER framework. All the files for the parts mentioned below are available on the SyntheCell GitHub.

---

**Protective box**

We designed a PMMA protective box to (1) keep the smart sleeve photodiodes from natural and artificial light to avoid interference with the OD measurements and (2) protect the electronic components in case a medium overflow occurs.

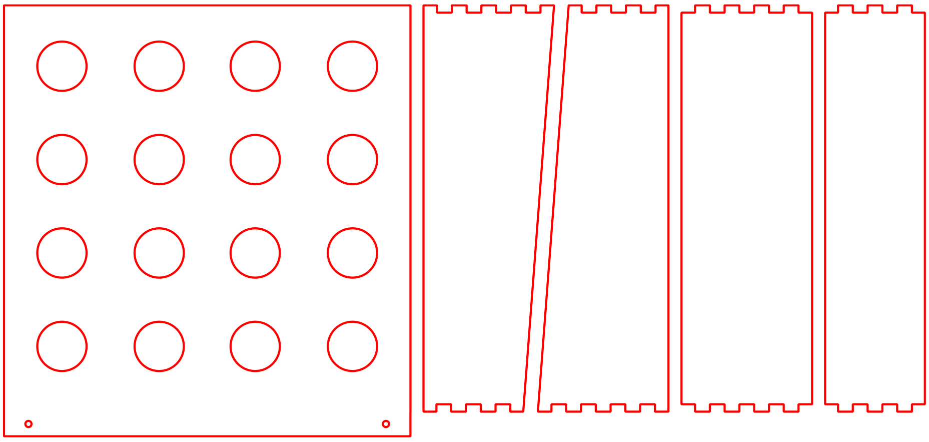

| Required parts | Required equipment |
|---|---|
| • 6 mm thick opaque PMMA | • $CO_2$ laser cutter |
| • Epoxy glue | Required blueprints |
| • Silicone glue | • *EVOBlackBox.pdf* |
| • 1 m of silicone tubing, outer diameter 6.4 mm | • *TopBarrierFront.pdf (x2)* |
| | • *TopBarrierSide.pdf (x2)* |

1. Cut the PMMA parts using a $CO_2$ laser cutter.
2. Using epoxy glue, glue the hinges of the front, sides and back walls together.
3. Glue the four top barriers to the top plate of the box.
4. Dry for a day.
5. Glue the top plate of the box on top of the walls and dry for a day.
6. Apply silicone glue to the top and the inside of the box to improve fixation and seal the box.
7. Cut two pieces of 50 cm silicone tubing. Connect the evacuation holes in the front of the box to a liquid waste container using the tubing.

---

**Neoprene pads**

Once the box above is positioned, the sleeve components must be isolated from potential liquid overflows. For this, square neoprene pads are positioned around the aluminum sleeves and brought into contact with the top of the box.

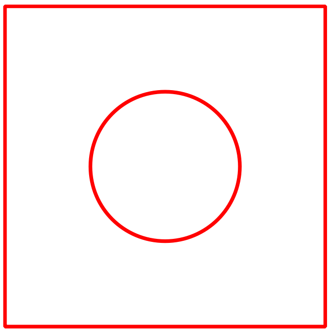

| Required parts | Required equipment |
|---|---|
| • 1.5 mm thick Neoprene rubber sheet | • $CO_2$ laser cutter |
| | Required blueprints |
| | • *NeoprenePad.pdf* |

1. Cut the Neoprene sheet using a $CO_2$ laser cutter.
2. Place the protective box on the eVOLVER.
3. Place the Neoprene pads around the aluminum sleeves.

---

**Cap silicone ring**

This silicone joint is placed on the inside of the vial cap to prevent medium from leaking through the cap thread in case of an overflow, as this may lead to liquid flooding the inside of the sleeve. The medium will then only leak through the sampling port, and protection from such issue is ensured by the new vial caps, protective eVOLVER box and neoprene pads above.

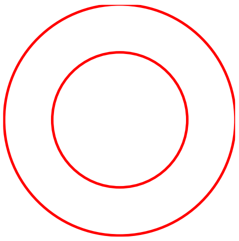

| Required parts | Required equipment |
|---|---|
| • 1/32” thick silicone rubber sheet | • $CO_2$ laser cutter |
| | Required blueprints |
| | • *CapRing.pdf* |

1. Cut the silicone sheet using a $CO_2$ laser cutter.
2. Place the silicone ring on the inside of the vial cap (see Section 3).

---

**Cap and cap holder**

To avoid movement of the vial throughout the experiment, we designed a set of a cap and a cap holder that fixes the vial inside the sleeve. These parts can be produced by 3D printing.

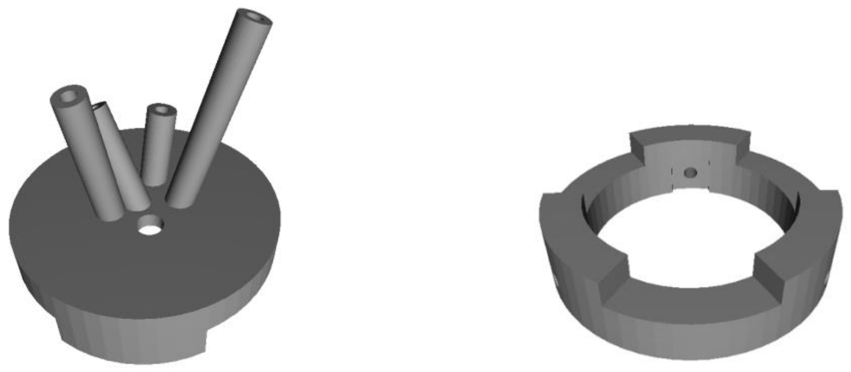

| Required parts | Required equipment |
|---|---|
| • Nylon PA11 | • 3D printer |
| | Required blueprints |
| | • *VialCap.stl* |
| | • *VialCapHolder.stl* |

---

**Switch protector**

To avoid accidentally switching off the eVOLVER system while manipulating the vials and fluidic parts, we designed a small protective box that hides the 5V and 12V switches behind a PMMA "door".

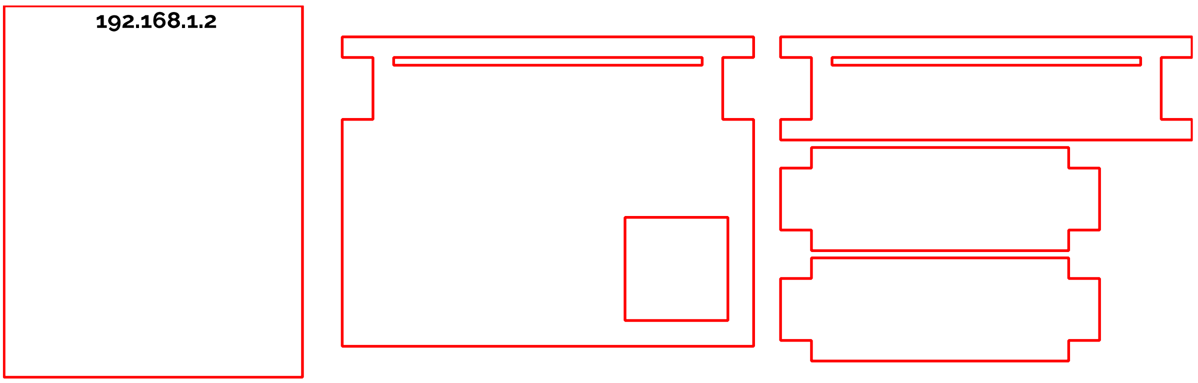

| Required parts | Required equipment |
|---|---|
| • 6 mm thick transparent PMMA | • $CO_2$ laser cutter |
| • 1 mm thick transparent PMMA | Required blueprints |
| • Epoxy glue | • *SwitchProtector.pdf* |

1. Cut the PMMA parts using a $CO_2$ laser cutter. For the protector "door", use 1 mm-thick PMMA.
2. Glue the base, sides, and top of the protector structure using epoxy glue.
3. Dry for one day.
4. Lift the eVOLVER and slide the part underneath to fit the front right rubber foot of the machine inside the hole in the base of the protector.
5. To protect the switches, slide the 1mm door through the gap.

#### 8.4. eVOLVER Telegram bot

We developed an alert system taking advantage of the open-source messaging application Telegram. It notifies the user in case of a syntax error in the code that would stall an experiment, or in case the experiment is paused for more than 10 min. This is extremely important, as a paused or stopped experiment will not dilute the cultures and these will reach saturation, with implications for the eVOLVER assay. Here we provide a step-by-step guide to configure this alert system for the eVOLVER framework.

1. Create a Telegram account.
2. Create a Telegram bot and obtain its access token. For this, follow the instructions in the section *"Obtain your bot token"* in this page.
3. Go to *web.telegram.org* and create a new channel. Set the visibility to *Private*.
4. Access the channel chat and identify the channel number in the URL.
5. The channel ID will be -100 followed by the number on the URL. Example:
   - URL = https://web.telegram.org/z/#-1234567890
   - ID = -1001234567890
6. Configure the Telegram bot by adding the token and channel ID in the *creds.json* file within the experiment folder (see Section 2.3).
7. Any user that joins the channel will receive alerts from eVOLVER.

#### 8.5. Components

This section provides a list of the specific components that we used to improve the eVOLVER framework for long-term experimental evolution with fission yeast.

**Bottles**

- Media bottles: Borosilicate bottles 1 L / 2 L
- Regular tubing: Silicone tubing ID 3.2 mm, OD 6.4 mm
- Tubing compatible with small hydrophobic molecules: Teflon (PTFE) tubing ID 3 mm, OD 6 mm
- Luer connectors adapted to the inner diameter of the bottle tubing

---

**Vials**

- Glass vials: Borosilicate 40 mL clear glass vials, 28x95cm, GPI 24/400
- Influx needle: 18G, 40 mm
- Efflux needle: 16G, 100 mm (adapt length to experiment requirements)
- Sampling needle: 20G, 70 mm
- Heat-shrink tubing: 6 mm, 2:1
- Silicone ring: Silicone rubber sheet, 0.8 mm thickness
- Magnetic stirring bar (Fisherbrand; Product code – 11818862)

---

**Vial caps**

- Nylon cap: 3D printed with Nylon PA11
- Screws: M3 x 4 mm headless flat end screws DIN 913
- EVA foam, 1 mm thickness

---

**Protective system**

- 6 mm thick opaque PMMA
- 6 mm thick transparent PMMA
- 1 mm thick transparent PMMA
- Neoprene pads: Neoprene rubber sheet, 1.5 mm thickness

---

**Miscellaneous**

- Spray glue for pumps: 3M™ Hi-Strength 90 Spray Adhesive
- Epoxy glue
- Silicone glue

**Complementary sources of information**

Although this document represents a standalone guide for using the improved eVOLVER system for long-term experiments, other applications may benefit from all the upgrades we have implemented. Nevertheless, additional knowledge of eVOLVER's functioning principles may be interesting for the user. The community of eVOLVER users is very active, and various inputs can be found at the following addresses:

eVOLVER forum: https://evolver.bio

eVOLVER Wiki: https://khalil-lab.gitbook.io/evolver/

Manufacturer's website: https://www.fynchbio.com/

Manufacturer's code repositories: https://github.com/FYNCH-BIO/

---

### LET'S EVOLVE !

---
